# Peptide Amphiphiles Hitchhike on Endogenous Biomolecules for Enhanced Cancer Imaging and Therapy

**DOI:** 10.1101/2024.02.21.580762

**Authors:** Li Xiang, Morgan R. Stewart, Kailin Mooney, Mingchong Dai, Samuel Drennan, Samantha Holland, Arnaud Quentel, Sinan Sabuncu, Benjamin R. Kingston, Isabel Dengos, Karla Bonic, Florian Goncalves, Xin Yi, Srivathsan Ranganathan, Bruce P. Branchaud, Leslie L. Muldoon, Ramon F. Barajas, Jared M. Fischer, Adem Yildirim

**Author notes:** These authors contributed equally: Li Xiang, Morgan R. Stewart, and Kailin Mooney.

## Abstract

The interactions of nanomaterials with biomolecules in vivo determine their biological fate. Here, we show that self-assembled peptide amphiphile (PA) nanostructures can dynamically interact with endogenous biomolecules and take advantage of naturally occurring processes to target a broad range of solid tumors. In circulation, self-assembled PA nanostructures disassemble and reassemble mainly with lipoproteins, which prolongs blood circulation and dramatically improves tumor accumulation and retention. Mechanistic studies suggested that PAs internalize into cancer cells by assembling with their cell membranes and independently of specific receptors. By exploiting these interactions, a PA developed in this study (namely SA-E) demonstrated specific accumulation in various xenograft, syngeneic, patient-derived xenograft, or transgenic rodent models. In addition, SA-E enabled the effective delivery of highly potent chemotherapy to different syngeneic and xenografted tumors with reduced side effects. With its simple and modular design and universal tumor accumulation mechanism, SA-E represents a promising platform for broad applications in cancer imaging and therapy.

## 1. Introduction

Peptide amphiphiles (PAs) spontaneously self-assemble into nanostructures in aqueous solutions through hydrophobic and electrostatic interactions and hydrogen bonding.^1–4^ By exploiting the rich chemistry of amino acid side chains and abundant peptide modifications at both the side chains and terminal ends, PA nanostructures with various morphologies have been developed.^5–7^ PA nanostructures are naturally biodegradable and biocompatible with low immunogenicity, making them promising delivery platforms for cancer therapy and detection.^8–11^ Like any other self-assembled structure, when PA nanostructures are introduced into circulation, they can disassemble as a result of the binding of their amphiphilic building blocks to biomolecules such as albumin or lipoproteins (LPs).^12,13^ Thus, the primary focus in the PA research has been to develop new nanostructures with good stability in the blood to allow them to carry payloads to their intended targets.^12–16^ While recent studies demonstrated the importance of bio-nano interactions on the biodistribution of nanomaterials, these interactions remained largely unexplored for PA nanostructures.^17–19^

In this work, we explored how PAs interact with endogenous biomolecules and cells and how these interactions affect their cancer targeting ability and biodistribution using various in vitro and rodent models. Our studies revealed a mechanism allowing PAs to target and enter cancer cells independent of cell state or surface markers. Specifically, we found that weakly assembled PAs can dynamically interact with blood biomolecules and cell membranes upon in vivo administration, allowing them to exploit the increased lipid metabolism of cancer cells to specifically accumulate in solid tumors. For these studies, we designed PAs that can self-assemble into micelles with spherical or rod-shaped morphologies, giving them different stabilities in blood plasma. Through molecular dynamic simulations and in vitro experiments, we found that intramolecular interactions in nanostructures of PAs with spherical morphology (namely SA-E) are weaker than rod-shaped ones (namely SA-K). Upon incubation with blood plasma, while more strongly interacting SA-K micelles mainly remained intact, SA-E micelles quickly disassembled due to lower intermolecular interactions in these structures. Protein chromatography and mass spectrometry studies showed that SA-E selectively binds to lipoproteins (LPs) in the blood, which are endogenous nanoparticles with sizes ∼5-80 nm (excluding chylomicrons) and composed of lipids, cholesterol, and amphiphilic proteins (i.e., apolipoproteins). LPs have long-blood circulation half-life, intrinsic biocompatibility (i.e., non-toxic and non-immunogenic), and tumor accumulation as a result of increased lipid metabolism of cancer cells.^20–23^ In addition, our studies indicated that PAs internalize into cells as monomers through membrane binding, not as intact micelles. Dynamic interactions of SA-E with these endogenous biomolecules prolonged its blood circulation and enabled strong accumulation in a broad range of solid tumors compared to more stable SA-K nanostructures. In addition, we found that SA-E was enriched in cancer cells in the tumor microenvironment. Biodistribution studies showed that SA-E was mostly cleared from normal tissue in 2 days but retained in the tumor for more than 2 weeks. The substantial tumor accumulation and retention of SA-E enabled the detection of millimeter-sized early breast tumors and colon adenomas in mice and intracerebral glioma tumors in rats with tumor-to-healthy brain tissue ratios of ∼16. Finally, we conjugated SA-E with a highly potent and toxic chemotherapeutic agent, Monomethyl auristatin E (MMAE), and demonstrated its strong antitumor efficacy in breast, glioma, and colon cancer models in mice with reduced side effects compared to the free drug.

## 2. Results and Discussion

### 2.1. Stability and interactions of PA nanostructures in blood plasma

The overall design of the PAs used in this study is shown in Figure 1a. Both PAs share a common motif to promote self-assembly through hydrophobic interactions and hydrogen bonding: GGGHAANG with a palmitic acid modification on the N-terminal.^24,25^ While keeping this motif the same, we tuned the hydrophilic part of the PAs to change the morphology of self-assembled PA nanostructures.^26^ SA-E and SA-K contain three glutamic acids (E) or lysines (K) in their hydrophilic part, respectively. In addition, a cysteine residue was added for dye or drug conjugation through maleimide-thiol coupling. PAs were conjugated with a near-infrared (NIR) fluorescent dye (indocyanine green; ICG) to track the PA nanostructures in vivo and investigate their stability in biological solutions. PA-ICG conjugates were purified using high-performance liquid chromatography (HPLC) and characterized using liquid chromatography-mass spectrometry (LC-MS) (Supporting Information, Figure S1,2).

**Figure 1.**
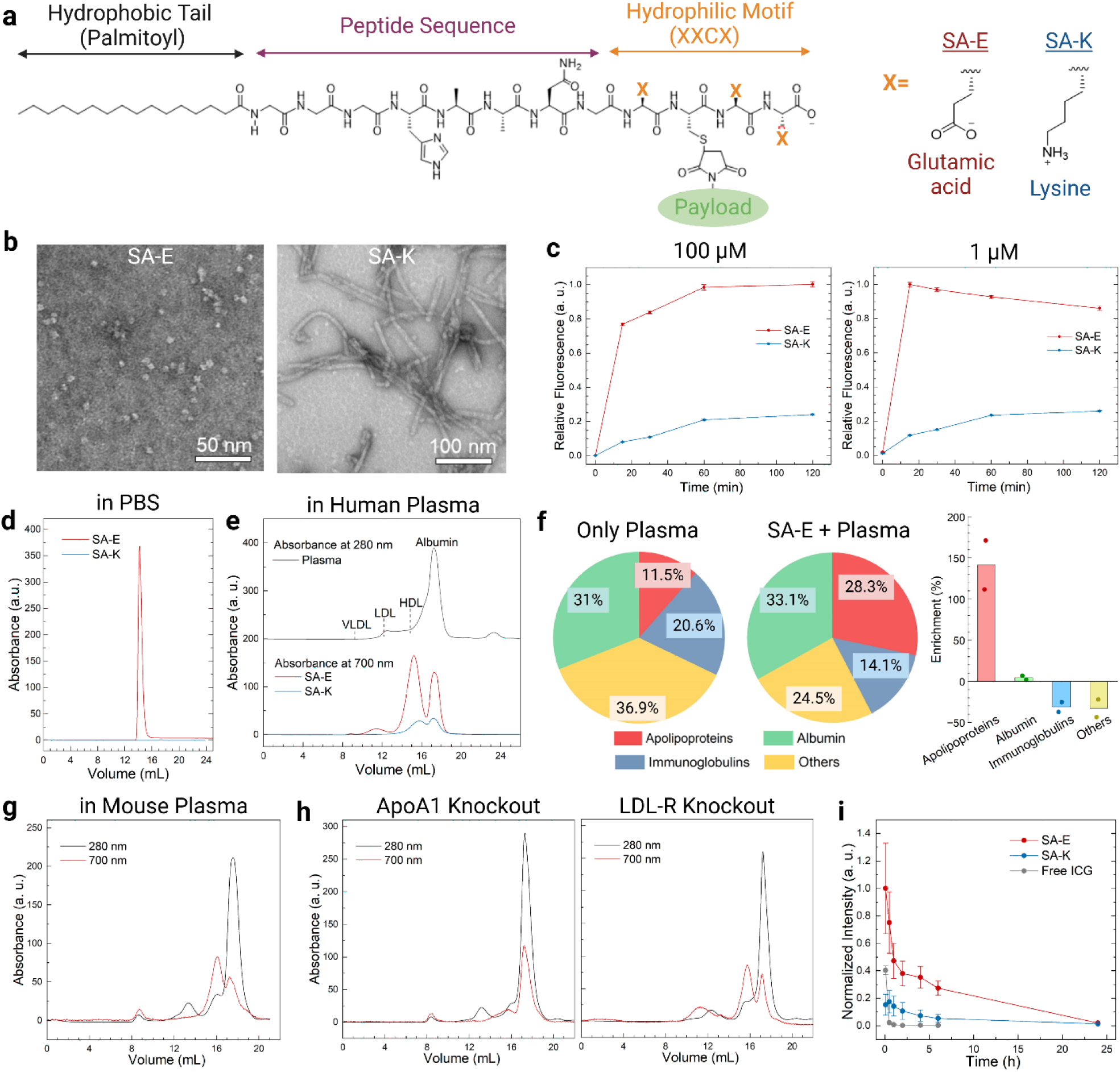
Spherical PA micelles disassemble in plasma and in situ reassemble with plasma biomolecules for prolonged blood circulation. a) Schematic showing the overall molecular structure of the PAs used in this study. b) TEM images of PA nanostructures. Samples were negatively stained using uranyl acetate. c) Fluorescence intensity of ICG labeled PAs in 10% human plasma over 2 hours. FPLC traces of PAs in PBS (d) or 10% human plasma (e). Absorbance at 280 nm was used to detect LPs and albumin, and 700 nm was used to detect SA-E. f) Summary of protein classes identified in pulled-down proteins in 10% human plasma samples with or without SA-E by mass spectrometry (left panel). Percent enrichment of proteins in the presence of SA-E (right panel). FPLC traces of plasma samples collected from wild type (g) or ApoA1-/-or LDL-R-/- (h) mice injected with SA-E (50 nmole) 1 hour before blood collection. i) Blood circulation of intravenously injected ICG labeled PAs or free ICG (50 nmole) in wild type mice. Data in (c, i) are presented as mean ± standard error of the mean (SEM). Bars in (f) are the mean of the data. Studies were run in triplicates in (c, i) and duplicates in (e).

The morphology of self-assembled PA nanostructures dispersed in phosphate-buffered saline (PBS, 10 mM, pH 7.4) was investigated using transmission electron microscopy (TEM) (Figure 1b). SA-E demonstrated a spherical micelle morphology with sizes around 5-10 nm, and SA-K formed rod-shaped micelles with diameters around ∼10 nm and lengths reaching several hundred nm.

To understand the stability of self-assembled PA nanostructures in biologically complex solutions, we initially utilized human plasma (Figure 1c). PAs at concentrations of 1 or 100 µM were incubated in 10% human plasma and fluorescence of ICG was measured at different time points. At both concentrations, the fluorescence of ICG was quenched in PBS in PA micelles due to the close packing of ICG molecules.^27^ While the addition of plasma resulted in an increase in the fluorescence of both PAs, suggesting partial or complete disassembly of their self-assembled nanostructures, the increase in the SA-E fluorescence was ∼3-fold larger and more rapid than SA-K at both concentrations. To gain more insights on the stability of PAs, we performed molecular dynamic simulations to study the intramolecular interaction in self-assembled PA nanostructures. In accordance with experimental results, simulations suggested more strongly assembled structures for the rod-shaped SA-K assemblies (See Supporting Information for details, Figure S3). Overall, these results showed that rod-shaped PA nanostructures of SA-K are more stable in plasma than spherical ones of SA-E.^28,29^

Next, we sought to identify the blood components that PA nanostructures interact with. The most likely interactions of PAs in the blood are expected to be with albumin and LPs (e.g., very low-density lipoprotein; VLDL, low-density lipoprotein; LDL, or high-density lipoprotein; HDL), as they are highly abundant in blood and carry lipophilic molecules.^20–23,30–33^ To study the interactions of PAs with these fractions of plasma, we performed fast protein liquid chromatography (FPLC) experiments using SA-E and SA-K in the absence or presence of pooled human plasma or purified LPs and albumin. Without plasma, SA-E dispersed in PBS eluted from the column as a single narrow peak, as expected for self-assembled micelles with a narrow size distribution of around 10 nm (Figure 1d). No peak was detected for SA-K suggesting that long rod-shaped micelles of SA-K could not pass the FPLC column. Before moving to the studies with human plasma, we determined the elution times of albumin and LPs (VLDL, LDL, or HDL) in the FPLC using human serum albumin or LPs purified from human plasma (Supporting Information, Figure S4a). In addition, we performed FPLC using SA-E mixed with purified LPs or albumin, which showed a lack of the SA-E micelle peak observed in PBS. SA-E eluted with the LPs or albumin (Supporting Information, Figure S4a), showing that SA-E micelles completely disassembled and reassembled with these biomolecules. Similarly, in 10% human plasma (Figure 1e) there was no intact SA-E micelle peak. Instead, SA-E was found to be mainly bound to LPs (65.3% total, 59.8%, 5.3%, and 0.3% for HDL, LDL, and VLDL, respectively) and to a lower extent to albumin (34.7%). While albumin peak in the FPLC trace at 280 nm was ∼20-fold stronger than HDL, SA-E bound to albumin (trace at 700 nm) was ∼2-fold lower than HDL, suggesting a stronger affinity of SA-E against HDL than albumin. The higher affinity of SA-E toward LPs was also confirmed by performing FPLC of SA-E incubated in a mixture of LPs and albumin, which showed almost entire binding to LPs (Supporting Information, Figure 4b). While a similar binding profile was observed for SA-K in human plasma (Figure 1e) with higher binding to LPs (55% total, 50.7% and 4.3% for HDL and LDL, respectively) than albumin (45%), the area under the curve (AUC) of all peaks was ∼4.5 lower than SA-E, which is in accordance with stability experiments above and further indicating that SA-K micelles remain mostly structurally stable in plasma. In addition, we prepared a version of SA-E without the n-terminal lipid modification and conjugated with ICG (No-SA, Supporting Information, Figure S5) to explore the impact of lipid modification on the assembly of PAs to plasma components. FPLC of No-SA in 10% human plasma (Supporting Information, Figure S6) showed that it remained mostly unbound in plasma (95.3%) with some binding to albumin (4.2%) and at a lower extent to HDL (0.5%), indicating that PAs bind to plasma components through their lipid modifications.

To further study the plasma components that PAs bind, mass spectrometry (MS) was used. For MS experiments, we only used SA-E due to its effective assembly with plasma components. A biotinylated version of ICG labeled SA-E was prepared (SA-Eb, Supporting Information, Figure 7) and incubated in human plasma. SA-Eb and the plasma components bound to it were pulled down using streptavidin-coated magnetic beads, and eluted proteins were analyzed using MS (Figure 1f). There were ∼2.4-fold more MS counts for the SA-Eb sample compared to proteins eluted from beads in the absence of SA-Eb (only plasma), suggesting that most proteins were specifically pulled down with SA-Eb (Supporting Information, Table S1). For the SA-Eb sample, 28.5% of all detected proteins were apolipoproteins (protein components of LPs), whereas the apolipoprotein content was 11.8% in the plasma only sample (Figure 1f and Supporting Information, Table S1). Apolipoprotein B, the main protein component of LDL and VLDL,^34^ was non-detectable in the plasma only sample, but it was abundant (5.4%) in the presence of SA-Eb. In addition, the concentration of Apolipoprotein A-I, the main protein component of HDL,^34^ increased ∼2.5-fold (to 18.4% from 7.3%) in the SA-Eb sample. The increase in the albumin concentration (to 33.3% from 31.8%) was lower than the increase in LPs, which is in accordance with FPLC results. Overall, these results further suggested that SA-E preferentially assembles with LPs in the blood.

### 2.2. Assembly of PAs with endogenous blood biomolecules prolongs circulation

We next explored the interactions of PAs with blood components in vivo to confirm that they interact with LPs in situ upon systemic delivery. For these experiments, we used SA-E due to its better binding ability to plasma components. ICG conjugated SA-E was intravenously injected into wild type mice, and blood samples were collected 1 h after SA-E injection. FPLC of plasma (Figure 1g) showed that most of SA-E was most preferentially bound to HDL again with similar percentages observed in human plasma (LPs: 65.8% total, 55.2%, 4.6%, and 5.9% for HDL, LDL, and VLDL, respectively and albumin: 34.2%). We also measured the ICG fluorescence in fractions of collected blood samples: plasma, white blood cells (WBC), and red blood cells (RBCs), to understand if SA-E also assembles or taken up by blood cells (Supporting Information, Figure 8). On average 98.9% of SA-E fluorescence was detected in the plasma with only 1.1% fluorescence in the RBC component suggesting that SA-E almost completely assembles with plasma components in the blood and its uptake by blood cells is negligible.

We then utilized two genetically engineered mouse models with aberrant lipid metabolism, ApoA1 and LDL-receptor (LDL-R) knockout mice (ApoA1-/- and LDL-R-/-), to further investigate LP specific binding of SA-E in blood and understand how the altered lipid metabolism affects interactions of SA-E with LPs. As SA-E mainly binds to HDL in the blood, we first used ApoA1-/-mice, which have significantly reduced levels of HDL due to the lack of the main apolipoprotein component of HDL in these mice.^35^ FPLC showed a dramatically reduced HDL binding (16.7%) in ApoA1-/-mice compared to wild type mice with significantly increased albumin binding (75.4%), confirming that SA-E mainly assembles with HDL in circulation (Figure 1h). We then utilized LDL-R-/-mice, which has elevated blood cholesterol and LPs, especially LDL,^36^ to explore how increased cholesterol levels affect the interactions of SA-E with plasma components. In general, a similar binding profile with wild type mice was observed for LDL-R-/-mice (Figure 1h) with slightly increased overall LP binding from 65.8% to 73%. A more significant difference was the increase in SA-E assembly with LDL and VLDL. While the LDL and VLDL were not separated as observed for wild type mice, total binding to these LPs increased to 28.6% from 10.6%. Nevertheless, SA-E was found to be mostly assembled with HDL (44.4%), showing that even a dramatic change in cholesterol levels does not significantly affect SA-E assembly with blood components.

After confirming the preferential assembly of PAs with LPs, especially HDL, we investigated how LP binding affects the blood circulation of PAs in wild-type mice. SA-E showed prolonged blood circulation compared to both SA-K and free ICG with a half-life of ∼1 hour (Figure 1i). In addition, the fluorescence of SA-E was 27% of initial intensity at 6 hours after injection, and it was still detectable 24 hours after injection (∼2% of initial fluorescence). For both SA-K and free ICG, a blood circulation half-live <5 min was observed. While SA-K fluorescence was still detectable 24 hours after injection, its blood concentration was lower (up to ∼6-fold) at all time points. In addition, the AUC of blood circulation plots was 4.3-fold higher for SA-E than SA-K. Blood concentration of free ICG was quickly decreased below 1% of the injected dose in an hour and it was not detectable 6 hours after injection, which is in accordance with the previous literature.^37^ We also studied the blood circulation of ICG labeled No-SA to further explore the effect of LP binding on blood circulation (Supporting Information, Figure S9). While No-SA showed improved blood circulation compared to free ICG, it was cleared more rapidly than SA-E with a half-life <1 hour and its fluorescence intensity decreased to around 5% of initial intensity 6 hours after injection. Altogether, these in vitro and in vivo experiments showed that SA-E nanostructures can effectively disassemble and reassemble with endogenous LPs in circulation, which prolongs blood circulation compared to structurally more stable assemblies of SA-K or non-LP binding No-SA.

### 2.3. Cellular entry of PAs

Next, we investigated interactions of PAs with cell membranes and the impact of LPs on these interactions using Cy5 labeled PAs (Supporting Information, Figure S10,S11). We first incubated PAs briefly (∼1 min) with 4T1 mouse breast cancer cells and imaged the cells at different time points after removing the PAs or no-SA. For both SA-E and SA-K, a strong Cy5 fluorescence was observed at the cell membrane shortly after adding the PAs (Figure 2a). Further incubation resulted in the internalization of PAs into cells. We again used No-SA (Cy5 labeled, Supporting Information, Figure S12) to explore the effect of lipid modification on cellular entry of PAs. No significant membrane binding or internalization was observed for No-SA (Supporting Information, Figure S13), indicating that PAs bind to cell membranes through their lipid modifications. While both SA-E and SA-K were effectively taken up by cells, SA-K showed ∼3-fold higher uptake than SA-E (Figure 2b), which might be due to its positive charge.^41^ These results also showed that PAs internalized in cells as monomers rather than taken up as intact micelles, which is in accordance with a previous report.^39-41^ To understand if PAs enter cells through a passive diffusion or endosomal pathways, we incubated cells with SA-E and an endosome stain (FITC-dextran) for 4 hours. Fluorescence of SA-E and endosomal fluorescein showed a good correlation with a Pearson coefficient of 0.605 ± 0.02 (Fig 2c, see also Video S1), suggesting that SA-E was internalized in cells mainly via endosomal uptake.

**Figure 2.**
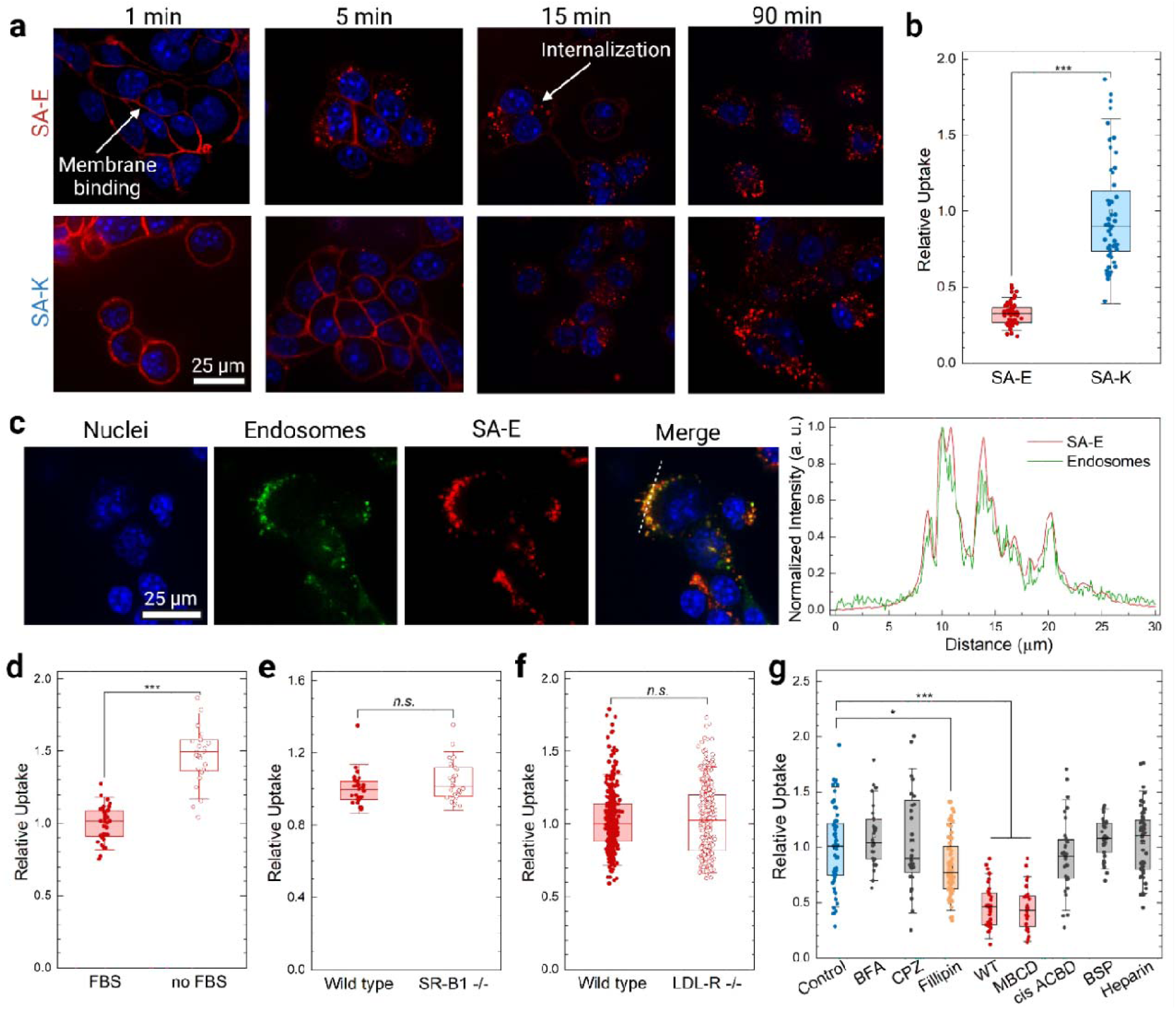
PAs internalize into cells through binding to lipid raft domains of cell membranes. a) Confocal microscope images of 4T1 cells incubated with SA-E or SA-K at different time points. b) Uptake of PAs by 4T1 cells. c) Confocal images of 4T1 cells incubated with endosomal stain (FITC-dextran) and Cy5 labeled SA-E. The panel on the right shows the intensity profile of FITC and Cy5 through the dashed line in the merged image. d) Uptake of SA-E by 4T1 cells in the presence or absence of FBS. e) Uptake of SA-E by wild type or SR-B1 knock out TRAMP-C2 mouse prostate cells. f) Uptake of SA-E by wild type or LDL-R knock out HeLa human cervical cancer cells. g) Uptake of SA-E by 4T1 cells in the presence of different inhibitors. BFA is Brefeldin A, CPZ is Chlorpromazine, WT is Wortmannin, MBCD is Methyl-b-cyclodextrin, and BSP is Bromosulfalein. The whisker box plots in (b,d-g) show the 25th and 75th percentiles with the median represented by the center line. Whiskers extend to 1.5x the inter-quartile ranges (IQR). Statistical analysis was performed using one-way analysis of variance (ANOVA) in (g) and Student’s t test in (b,d-f). *n.s*. is non-significant, \**p* <0.05 and \*\*\**p* <0.001.

Next, we explored the effect of LPs on the cellular uptake of PAs using SA-E. First, we studied the uptake of SA-E by 4T1 cells in the presence or absence of blood biomolecules by using cell media with or without fetal bovine serum (FBS). The absence of FBS in cell media did not decrease SA-E uptake. On the contrary, SA-E uptake was ∼1.6 higher in FBS-free media (Figure 2d), which suggests a competitive binding of SA-E with blood biomolecules (mainly LPs) and cell membranes that can decrease the uptake. In addition, we studied the impact of two well-known LP receptors for LDL and HDL, LDL-R and Scavenger receptor B type 1 (SR-B1), respectively,^20,42^ on SA-E uptake using cells knockout for these receptors. We did not observe any significant change in the cellular uptake of SA-E in LDL-R knockout (LDL-R-/-) HeLa cervical or SR-B1 knockout (SR-B1-/-) TRAMP-C2 mouse prostate cancer cells compared to wild type controls (Figure 2e,f).^43^

To further evaluate the uptake mechanism of SA-E, we used a series of inhibitors blocking different endocytosis pathways or cell surface receptors (Figure 2g).^44-47^ Significant decreases in the cellular uptake were observed for Wortmannin (WT, a PI3K inhibitor and general inhibitor of endocytosis) and Methyl-b-cyclodextrin (MBCD, extracts cholesterol from cell membranes). In addition, Fillipin, a caveolae-dependent endocytosis and a binder of membrane cholesterol,^48^ decreased SA-E uptake to a lesser extent. As cholesterol is enriched in lipid rafts, which are solid domains of cell membranes, ^49-51^ these results indicate a higher affinity of SA-E toward cholesterol-rich lipid-raft domains of cell membranes. Since SA-E contains several anionic residues, we also used inhibitors of glutamate uptake (cis-ACBD) or organic anionic uptake (bromosulfalein, BSP)^52,53^ and found no change in SA-E uptake. Altogether, these results suggest that PAs are internalized in cancer cells via membrane binding, especially to cholesterol-rich domains, independent of LP or other cell surface receptors.

### 2.4. Dynamic interactions of PAs with endogenous biomolecules improve tumor accumulation

Next, we studied the biodistribution of PAs in tumor-bearing mice to understand how their interactions with plasma components and cell membranes affect their accumulation in solid tumors and healthy tissues. As discussed above, intact micelles of PAs are not fluorescent. Thus, fluorescence imaging can only detect disassembled PAs. Therefore, we first investigated the disassembly of PAs in vivo by incubating them in mouse liver homogenates. SA-E and SA-K showed almost identical fluorescence intensity in liver homogenates (Supporting Information, Figure S14) indicating a similar disassembly degree for SA-E and SA-K in the tissues and confirming that fluorescence imaging using an in vivo imaging system (IVIS) could be used to compare biodistribution of these two PAs with different plasma stability.

We initially evaluated the tumor accumulation of ICG conjugated SA-E and SA-K (50 nmole, intravenously injected) within a syngeneic mouse 4T1 breast cancer model. We also used free ICG injection as a control. IVIS imaging showed a significantly higher tumor accumulation for SA-E compared to both SA-K and free ICG (Figure 3a). For SA-E, maximum fluorescence levels were detected in the tumor at 11 hours post-injection (Figure 3b). SA-E signal slowly decreased over 2 weeks with a 50% decrease in the signal on day 3 and still detectable levels even 2 weeks after injection demonstrating its prolonged retention. Background signal was mostly cleared 2 days after injection (Figure 3a), and tumor to background signal ratio (TBR) remained above 2.5 from 11 hours to 2 weeks post-injection with a maximum of around 5-6 from days 2 to 8 (Figure 3c). SA-K, with better stability in circulation, reached a maximum at 4 days post-injection and it was ∼6-fold lower than the maximum signal of SA-E. Also, the total tumor accumulation of SA-K was ∼2.7-fold lower than SA-E (Supporting Information, Figure S15). In addition, the maximum TBR was ∼2-fold lower than that of SA-E. Free ICG did not identify the tumor at any time point, and its TBR ratio always remained around 1. In addition, we evaluated the tumor accumulation of ICG conjugated No-SA to understand the effect of LP binding on cancer targeting (Supporting Information, Figure S16a,b). ICG conjugated to No-SA showed significantly improved tumor accumulation than free ICG, which can be explained by the improved blood circulation of No-SA. Compared to SA-E, No-SA accumulated more quickly in the tumor and mostly cleared in 2 days. Total tumor accumulation of SA-E was ∼1.6-fold higher than No-SA (Supporting Information, Figure S15). In addition, No-SA showed significantly lower TBR than SA-E (Supporting Information, Figure S16c).

**Figure 3.**
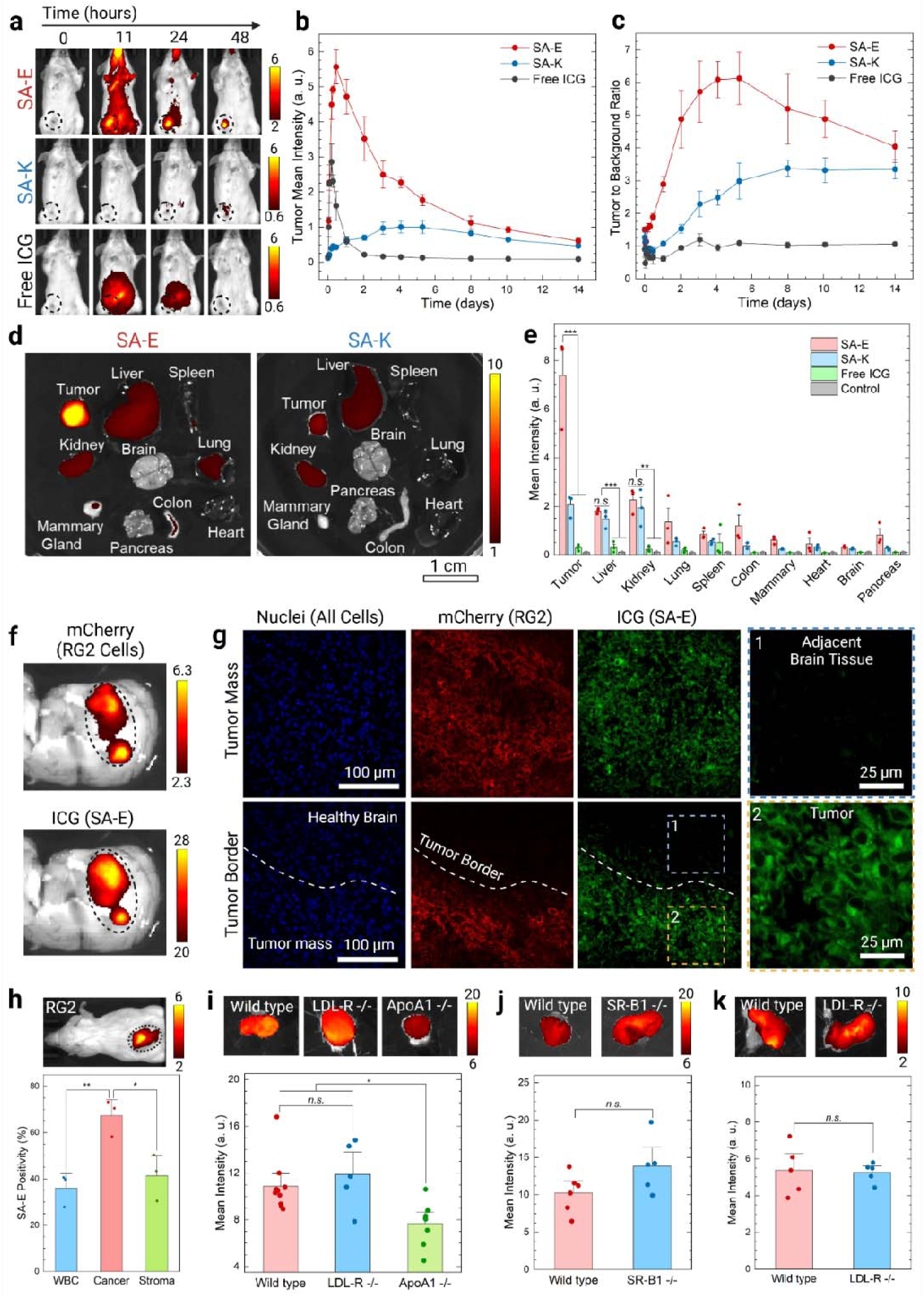
SA-E shows strong tumor accumulation and retention. a) Representative IVIS images of intravenously injected ICG labeled SA-E or SA-K (50 nmole) or free ICG in 4T1 tumor-bearing mice at different time points. Black circles highlight the tumor location. b) Calculated mean intensities of ICG signal and c) tumor to background signal ratio of SA-E, SA-K, and free ICG at different time points after injection. d and e) Biodistribution of SA-E, SA-K, or free ICG in 4T1 tumor-bearing mice. d) Representative IVIS images showing accumulation of SA-E and SA-K in tumors and major organs. Tissues were excised 2 days after injection. e) Mean ICG fluorescence intensity in the tumor and major organs of mice received SA-E, SA-K, or free ICG injection compared to control mice. f) IVIS imaging of mCherry expressing RG2 tumor sections showing a good overlap between the mCherry signal of RG2 cells and the ICG signal of SA-E. g) Confocal microscope images of a brain tissue section with mCherry expressing RG2 tumors showing specific accumulation of SA-E in RG2 tumors and internalization in cells in the tumor microenvironment. h) Flow cytometry analysis of mCherry expressing RG2 xenografts in mice that received SA-E injection (50 nmole) 2 days before harvesting tumors. The top panel is an IVIS image of RG2 tumor-bearing mice showing an accumulation of SA-E in the tumor. i) Mean tumor intensity of SA-E 2 days after injection (50 nmole) into MC-38 tumor-bearing wild type, LDL-R-/-, or ApoA1-/- mice. Images on the top are representative IVIS images of the tumors. j) Mean tumor intensity of SA-E 2 days after injection (50 nmole) into wild type or SR-B1-/- TRAMP-C2 tumor-bearing mice. Images on the top are representative IVIS images of the tumors. k) Mean tumor intensity of SA-E 2 days after injection (50 nmole) into wild type or LDL-R-/- HeLA tumor-bearing mice. Images on the top are representative IVIS images of the tumors. Data are presented as mean ± SEM. Studies were run in at least triplicates, except for ICG accumulation in the colon, mammary gland, heart, brain, and pancreas in (e) was obtained using a single mouse, and SA-K accumulation in the colon in (e) was obtained using two mice. Statistical analysis was performed using one-way analysis of variance (ANOVA) in (e, h, and i) and Student’s t test in (j and k). *n.s*. is non-significant, \**p* <0.05, \*\**p* <0.01, and \*\*\**p* <0.001.

Next, we investigated the biodistribution of PAs and free ICG in Balb/c mice bearing 4T1 tumors by imaging the organs 2 days post-injection (50 nmole). In line with live animal imaging, a significantly stronger fluorescence in the tumor was observed for SA-E compared to SA-K and free ICG (Figure 3d,e and Supporting Information, Figure S17). Notably, SA-E demonstrated a high tumor-to-liver signal ratio of 4.1, whereas it was 1.4 for SA-K. To estimate the percent injected dose (%ID) in each organ, tumors and major organs were homogenized and ICG fluorescence in each organ was measured (Supporting Information, Figure S18). In accordance with IVIS imaging, SA-E concentration in the tumor (1.9 ± 0.4 %ID / g tissue) was significantly higher than liver (0.8 ± 0.1 %ID / g tissue) and other major organs. Overall, these results suggest that SA-E and SA-K are mainly transported to the solid tumors by exploiting endogenous biomolecules, and better binding ability to plasma components improves biodistribution. Due to its better tumor accumulation and biodistribution, we used SA-E in the rest of the study.

We studied the clearance pathway and toxicity of SA-E in wild type mice. For clearance studies, SA-E (50 nmole) was injected into wild type mice, and urine and stool samples were collected at different time points (Supporting Information, Figure S19). Strong ICG signal was detected in the stool samples, which peaked at 6 hours after injection and mostly diminished at 2 days. No fluorescence signal was detected in urine samples at any time point. These results indicated that SA-E was cleared through liver and intestines as expected for LPs.^54^ The toxicity of SA-E was studied in wild type mice using a 4-fold higher dose than the optimal imaging dose (200 nmole). We measured complete blood counts (CBC) for blood toxicity, Aspartate transferase (AST) for liver toxicity, and Creatinine for kidney toxicity from blood samples collected 1 day after SA-E injection and found no difference between SA-E or control, suggesting SA-E is highly biocompatible (Supporting Information, Figure S20).

To further evaluate tumor-specific accumulation and in vivo cancer cell internalization of SA-E, we next utilized an orthotopic rat RG2 glioblastoma model. For these experiments, RG2 cells were injected into the rat cerebrum, and their growth was followed using gadolinium contrast-enhanced magnetic resonance imaging (MRI). ICG labeled SA-E (500 nmole) was injected when the tumors were easily visualized by MRI. Figure S21 (Supporting Information) shows the distribution of SA-E in the organs of a RG2 tumor-bearing rat, where significant ICG fluorescence was observed in the RG2 cell injection site with significantly lower accumulation in major organs. To better image the tumors, brains were wholemount bisected through the approximate middle of the tumor in a similar orientation as the MRI. There was near complete overlap between the T1 MRI signal and SA-E signal (Supporting Information, Figure S21b). Overall, the SA-E signal in the tumor was ∼16-fold higher than the adjacent brain tissue (Supporting Information, Figure S21c). To confirm the specific accumulation of SA-E in cancer cells, we used mCherry expressing RG2 cells. IVIS imaging of a wholemount bisected brain sample showed an almost complete overlap between the mCherry fluorescence of RG2 cells and ICG fluorescence of the SA-E (Figure 3f). Confocal microscopy showed cellular internalization of SA-E in RG2 cells and uniform accumulation in the tumor tissue with clear tumor margins and no detectable fluorescence in the healthy brain tissue (Figure 3g).

We used flow cytometry to explore if SA-E is taken up more by cancer cells compared to other cell types in the tumor microenvironment due to increased lipid uptake of cancer cells.^20^ For these studies, we used mCherry expressing rat glioblastoma (RG2) xenografts in mice, and Cy5 conjugated SA-E. We first confirmed the accumulation of SA-E in RG2 tumors using ICG labeled SA-E (Figure 3h). Then, Cy5 labeled SA-E was injected into tumor-bearing mice, tumors were harvested 2 days after SA-E (50 nmole) injection, and flow cytometry was performed for RG2 (mCherry), white blood, and other cells in the tumor stroma. mCherry cancer cells also containing Cy5 were found ∼70% of the time, whereas CD45 positive only (white blood cells) or CD45 and mCherry negative cells (tumor stroma) contained Cy5 ∼35-40% of the time (Figure 3h and Supporting Information, Figure S22). These results demonstrated that SA-E can effectively internalize in cancer cells and is enriched (∼2-fold) in cancer cells compared to other cells in the tumor environment.

Next, we performed in vivo experiments using different knockout strategies to study the proposed LP hitchhiking based tumor accumulation mechanism of PAs. First, we studied the tumor accumulation of SA-E in syngeneic MC38 colon cancer tumors in wild type, ApoA1-/-, or LDL-R-/- mice (Figure 3i). Compared to the tumors formed in wild type mice, SA-E uptake was significantly lower in ApoAI -/- mice, suggesting that assembly with HDL is critical for SA-E to accumulate in solid tumors. On the other hand, there was no significant change in LDL-R-/- mice, indicating that alterations in the cholesterol levels do not significantly affect the tumor targeting of SA-E. In addition, SA-E accumulation in other organs was not significantly different from wild type control for both models (Supporting Information, Figure S23). Next, we wanted to understand the role of receptor-mediated uptake in the cancer cells in vivo using SR-B1 knockout TRAMP-C2 mouse prostate cancer and LDL-R knockout human cervical cancer HeLa cells with their wild type controls (Figure 3j,k). In accordance with cell culture results, SR-B1 or LDL-R was not needed for tumor accumulation. Overall, these results suggest a mechanism of dynamic assembly to LPs in circulation followed by another dynamic assembly for a non-receptor mediated cancer cell uptake occurs in vivo and is supported by our initial in vitro experiments. The proposed mechanism for cancer-specific accumulation of PA nanostructures with low structural stability is summarized in Figure 4.

**Figure 4.**
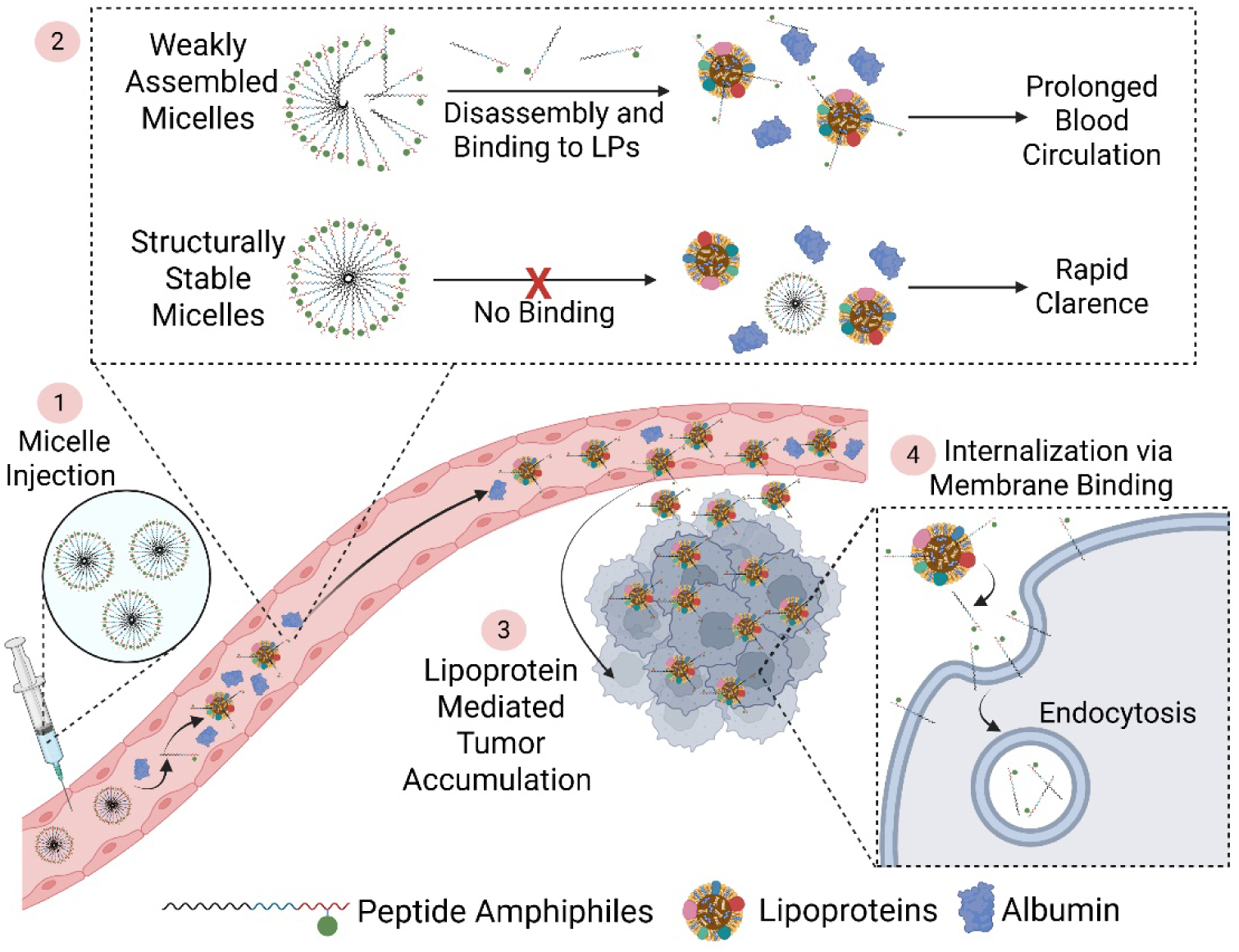
Schematic showing the proposed tumor accumulation and cancer cell internalization mechanisms of SA-E. Created with Biorender.com.

### 2.5. SA-E accumulated in a broad range of solid tumors

Since the proposed tumor-targeting mechanism of SA-E is universal, we tested tumor accumulation in a broad range of cell types and tissue locations. In addition to the 5 tumor models discussed above, we utilized 8 additional xenograft, syngeneic, transgenic, or PDX tumor models in mice. SA-E demonstrated highly specific tumor accumulation in all 13 tumor models of 8 different tissue types (Figure 4a). The tumors used in these studies ranged in size from a few millimeters to a centimeter in syngeneic models in immune competent mice as well as human or rat cells in immunocompromised mice. Strong tumor signal and high tumor-to-background ratios ranging between ∼3.5-50 were observed for the tumors imaged in live animals (Figure 4b). Also, by utilizing A375 human melanoma cell line expressing luciferase, we showed that the SA-E signal overlapped where the tumor was located. In addition, SA-E demonstrated strong accumulation in 2 transgenic models: MMTV-PyMT and APC^min^. MMTV-PyMT mice develop early to late cancer lesions in all mammary glands with time.^55^ Figure 4a shows representative images from an MMTV-PyMT mouse injected with SA-E weekly up to 85 days of age when all mammary glands developed tumors with different sizes. Notably, SA-E accumulated in early small lesions at 71 days of age, which developed large tumors 1-2 weeks later. In addition, organs harvested at the end of the experiment showed strong accumulation in mammary tumors but not in other organs (Supporting Information, Figure S24). APC^min^ mice is a transgenic colon cancer model suitable for studying early adenoma lesions.^56^ Small intestinal adenomas and colon polyps of APC^min^ mice injected with SA-E at 4 months of age had higher fluorescent signals than the surrounding normal intestine tissue (Figure 4a). Overlaying the identified tumors with SA-E fluorescence signal revealed that 100% of small intestinal adenomas and colon polyps were positive. In total, 71/71 small intestinal adenomas and 4/4 colon polyps were detected in 3 mice (Supporting Table 2) by marking all adenomas and polyps using the photograph images. In addition, a good linear correlation between lesion size and SA-E signal was observed with tumors even smaller than 1 mm^2^ being identified (Figure 4c). These results strongly suggest that SA-E uses a universal mechanism to target a broad range of tumors, including millimeter-sized, early tumor lesions.

### 2.6. SA-E enables more efficient and safer chemotherapy

Based on strong tumor accumulation and retention of SA-E, we hypothesized that SA-E can improve the efficacy of chemotherapies and decrease their side effects. To evaluate SA-E in drug delivery to solid tumors, we conjugated a highly potent tubulin inhibitor, Monomethyl auristatin E (MMAE),^57^ using an MMAE prodrug (vcMMAE)^57^ with a Cathepsin B cleavable linker (Supporting Information, Figure S25). SA-E-MMAE inhibited the growth of 4T1 and RG2 cells in vitro with IC50 values of 1.18 µM and 0.21 µM, respectively (Supporting Information, Figure S26).

We then tested SA-E-MMAE in the 4T1 orthotopic mouse model. 4T1 tumor bearing mice were intravenously injected with 5 doses of free MMAE (0.1 or 0.3 mg/kg) or SA-MMAE (0.3 or 0.6 mg/kg, based on MMAE) over 11 days. Free MMAE injections resulted in a slight reduction in tumor growth (Figure 6a). While free MMAE was tolerated at 0.1 mg/kg, increasing the dose to 0.3 mg/kg resulted in high toxicity with rapid weight loss, and treatment needed to be stopped after the third injection (Figure 6b). SA-E-MMAE was tolerated much better at higher doses. There was no significant weight loss at a 2-fold higher dose of 0.6 mg/kg with SA-E-MMAE. Importantly, SA-E-MMAE showed significant tumor growth reduction (∼75%) compared to both free MMAE and PBS control tumors at the dose of 0.6 mg/kg (Figure 6a). Next, we tested the antitumor efficacy of SA-E-MMAE against RG2 xenografts in nude mice. For this experiment, we further increased the SA-E-MMAE dose to 1 mg/kg. Similar to 4T1 results, we observed a significant reduction in the tumor volume, around 80%, compared to the control (Figure 6c). While a slight weight loss was observed at the beginning of the therapy, overall, the mice tolerated 5 doses of SA-E-MMAE injections at 1 mg/kg (Figure 6d). Finally, we tested the SA-E-MMAE in HCT-116 tumor bearing mice. We again applied 5 doses (0.8 mg/kg of MMAE), but this time we started the treatment when the tumor volume was around ∼50-100 mm^3^ on day 11 to investigate if SA-E-MMAE treatment can result in tumor shrinkage. All mice that received SA-E-MMAE showed reduction in the tumor size (Figure 6e), which significantly improved the overall survival (Figure 6f). Average tumor volume was ∼150 mm^3^ at the end of SA-E-MMAE treatment on day 21, which gradually decreased to around 40 mm^3^ on day 33 and remained almost unchanged for 2 weeks. While tumors in 3 of the mice started to grow again on days between ∼50 and 80, one mouse remained tumor-free on day 120. Overall, these results suggest that SA-E can be conjugated to chemotherapy drugs to improve their therapeutic index and antitumor activity.

**Figure 5.**
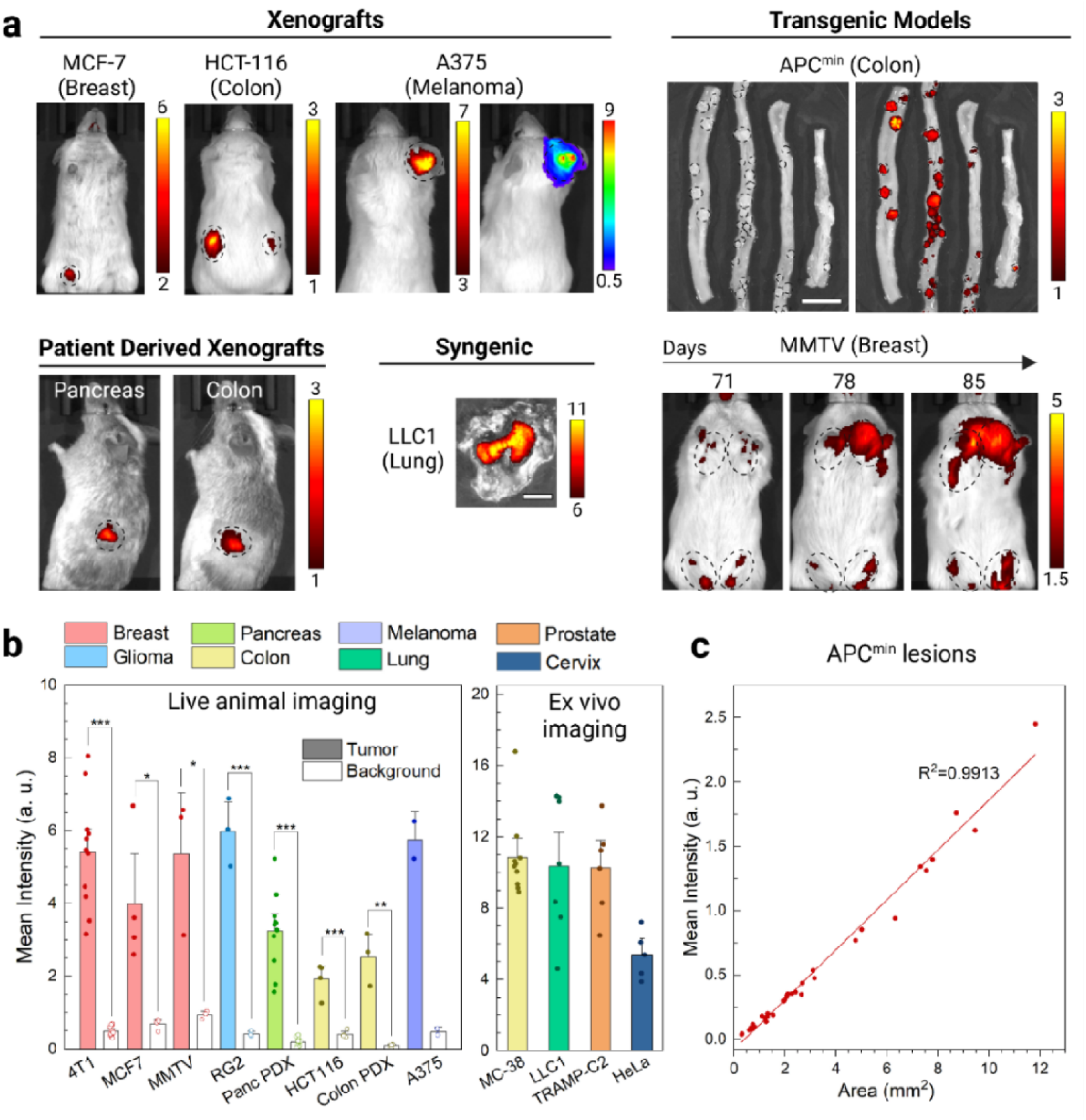
SA-E accumulates in a broad range of solid tumors. a) Representative IVIS images showing the SA-E accumulation in various tumor models of seven different tissues. MCF7, A375, and HCT-116 are human cell lines. LLC1 is a mouse cell line. In addition, two patient-derived xenografts (PDXs) of human colon and pancreatic cancer were used. MMTV and APC^min^ are transgenic mouse models of breast and colon cancer, respectively. For the MMTV model, mice were imaged weekly. For the APC^min^ model, bright field (left) and fluorescence (right) IVIS images show small intestine and colon samples of an APC^min^ mouse injected with SA-E at 4 months of age. IVIS imaging was performed 2 days after ICG labeled SA-E injection (50 nmole) either using live animals or excised tumors. For the luciferase-expressing A375 model, luciferase imaging was also performed (right panel), showing a good overlap between ICG and luciferase signals. Black circles highlight the areas with tumors, adenomas or polyps. Scale bars are 1 cm. b) Tumor and background intensity of SA-E in different models: live animal imaging (left) and ex vivo imaging (right). c) Correlation between lesion size and SA-E signal in the lesions detected in the small intestines and colons of three Apc^min^ mice. Data are presented as mean ± SEM. Studies were run in at least triplicates except the A375 model, which was performed in duplicate. Statistical analysis was performed using Student’s t test. \**p* <0.05, \*\**p* <0.01 and \*\*\**p* <0.001. No statistical analysis was performed for the A375 model.

**Figure 6.**
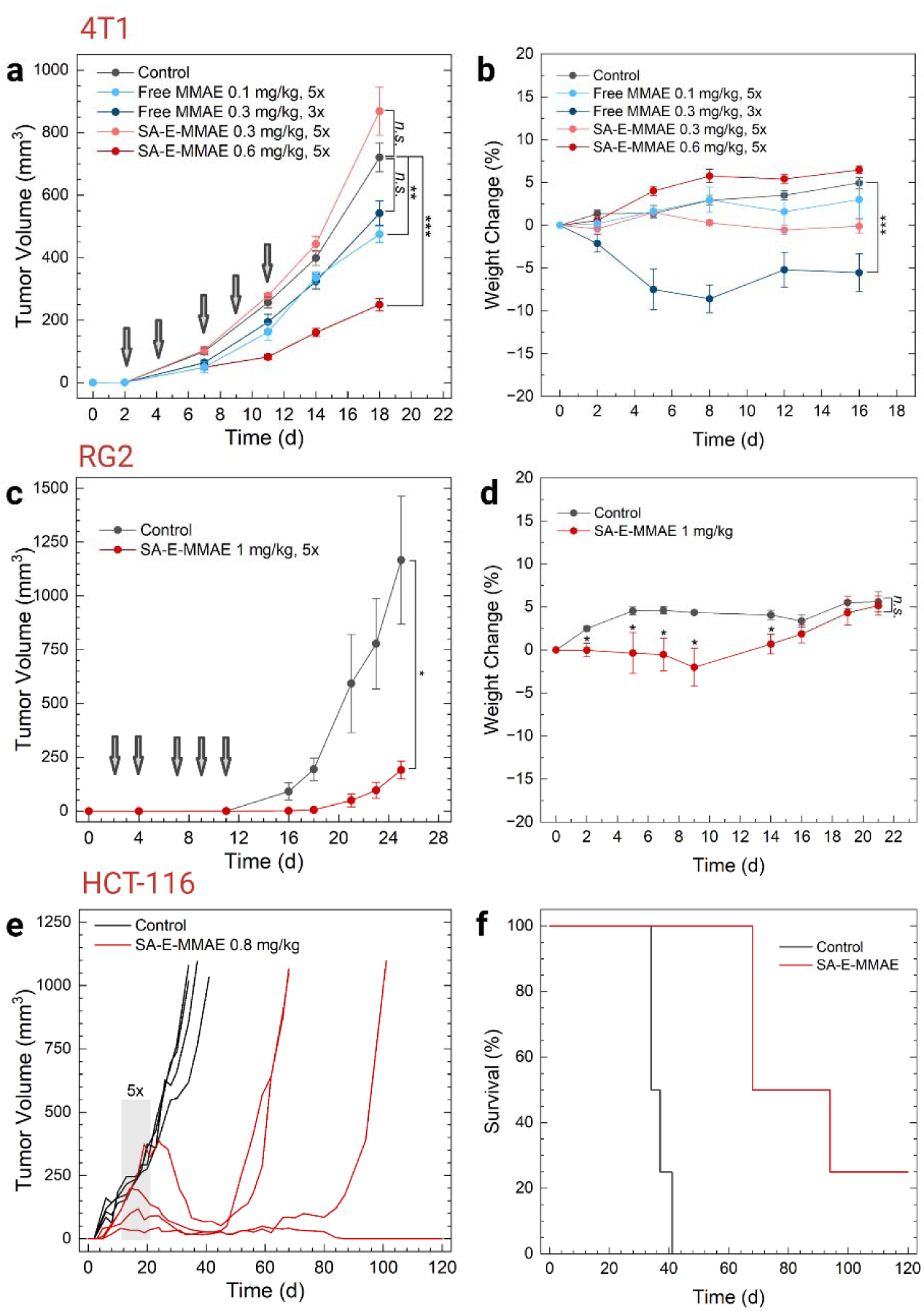
SA-E improves the antitumor efficacy and safety of MMAE. a) Tumor volume of 4T1 tumor bearing mice received PBS (n=18), 0.1 mg/kg of free MMAE (n=3), 0.3 mg/kg of free MMAE (n=5), 0.3 mg/kg of SA-E-MMAE (n=5), or 0.6 mg/kg of SA-E-MMAE (n=5). All mice received 5 doses of PBS, free MMAE, or SA-E-MMAE except 0.3 mg/kg free MMAE group, which received first 3 doses due to significant weight loss. b) Average body weight change of the mice that received the treatments in (a). c) Tumor volume of RG2 tumor bearing mice received 5 doses of intravenous injection of PBS (n=4) or 1 mg/kg of SA-E-MMAE (n=4). d) Average body weight change of the mice that received the treatments in (d). e) Tumor volume of HCT-116 tumor bearing mice received 5 doses of intravenous injection of PBS (n=4) or 0.8 mg/kg of SA-E-MMAE (n=4) on days 11, 14, 17, 19, and 21. f) Overall survival plot of the mice in (e). Grey arrows in (a) and (c) and grey box in (e) show the days that the mice received PBS or drug injections. Data are presented as mean ± SEM. Statistical analysis was performed using two-way analysis of variance (ANOVA) in (a) and (b) and Student’s t-test in (c) and (d). *n.s.* is non-significant, \**p* <0.05, \*\**p* <0.01, and \*\*\**p* <0.001.

## 3. Conclusions

In summary, we present a new mechanism for weakly assembled PA nanostructures (SA-E) to specifically accumulate in a broad range of solid tumors based on their in-situ assembly with plasma biomolecules and cell membranes. The proposed cancer targeting mechanism of SA-E is summarized in Figure 4. In circulation, SA-E disassembles and reassemblies mainly with LPs and to a lower extent with albumin (Figure 1c-h). Hitchhiking on plasma components provides prolonged blood circulation (Figure 1i), allowing SA-E to pass the tumor site numerous times and enabling their effective accumulation in the tumor microenvironment due to increased lipid uptake of solid tumors. In the tumor site, SA-E internalizes into cancer cells mainly through binding to cell membranes, especially to cholesterol-rich domains (Figure 2). Using these endogenous interactions, SA-E demonstrated high tumor accumulation and retention in various mouse and rat models, including transgenic models, patient-derived xenografts, orthotopic glioma tumors, and early lesions. In addition, we showed that SA-E can improve the efficacy of a chemotherapy drug with reduced side effects. The findings of this study underscore the importance of interactions of self-assembled nanomaterials with biomolecules on their fate in vivo and represent a novel way of exploiting these interactions to develop platforms with universal cancer-targeting ability.

## 4. Experimental Section

### Materials

Peptides were commercially obtained from Genscript with purity >95%. The following reagents were used: ICG maleimide (Iris Biotech, RL-2820), ICG (Fisher Scientific, I0535100MG), Cy5 maleimide (BroadPharm, BP-22552), vcMMAE (MedChem Express, HY-15575), MMAE (MedChem Express, HY-15162), HDL from human plasma (Sigma-Aldrich, SAE0054), LDL from human plasma (Sigma-Aldrich, SAE0053), VLDL from human plasma (Millipore Sigma, 437647), human serum albumin (Fisher Scientific, OSRHSA10), pooled human plasma (Innovative Research, IPLAWBK2E50ML), FITC-dextran 70 (Santa Cruz Biotechnology, sc-263323), brefeldin A (Alfa Aesar, J62340), bromsulphalein (MedChem Express, HY-D0217), chlorpromazine hydrochloride (Sigma-Aldrich, C8138), cis-ACBD (Tochris, 0271), filipin complex (Sigma-Aldrich, F9765), heparin sodium salt (Sigma-Aldrich, H3149-10KU), methyl-β-cyclodextrin (Sigma-Aldrich, C4555-5G), wortmannin (Alfa Aesar, J63983), bovine serum albumin (VWR, 0332), 4’,6-diamidino-2-phenylindole (DAPI) (Sigma-Aldrich, D9542), Hoechst stain (Invitrogen, H3570), 3-(4,5-dimethylthiazol-2-yl)-2,5-diphenyltetrazolium bromide (MTS) cell proliferation assay (Abcam, ab197010). Gadodiamide (GdDTPA-BMA, Omniscan®, GE Healthcare Inc. Marlborough, MA) MRI contrast was purchased through the OHSU in-patient pharmacy. Manufacturers and catalog numbers of additional materials, such as antibodies, are given in the text below.

### Preparation of PA dye or drug conjugates

4 mg of SA-E (2.92 μmol) and 3.21 μmol (1.1 eq) of maleimide-functionalized dyes or drugs were mixed in 2 mL of anhydrous DMSO (Sigma Aldrich, 900645). The mixture was incubated at room temperature overnight under gentle shaking (300 rpm). The same protocol was used to conjugate ICG or Cy5 to SA-Eb, SA-K or No-SA. The progression of the reaction was tracked using LC-MS, employing an Acquity UPLC system (Waters) equipped with a SQ Detector 2 (Waters) and a reverse-phase column (Waters, ACQUITY UPLC BEH C18 Column, 130Å, 1.7 µm, 2.1 mm x 50 mm). A solvent gradient from water/acetonitrile (95:5) with 0.1% formic acid to water/acetonitrile (5:95) was run over 9 minutes at a rate of 0.4 mL/min. Then, purification of the reaction mixtures was conducted using HPLC (Waters 1525, binary HPLC pump) with a reverse-phase column (Waters, XBridge BEH C18 OBD Prep Column, 130Å, 5 µm, 19 mm x 150 mm). The solvent gradient in the HPLC method was gradually adjusted from water/acetonitrile (95:5) to (5:95) with 0.1% trifluoroacetic acid over a 45-minute run at a flow rate of 0.8 mL/min. A UV/Visible detector (Waters 2489) was applied to monitor the absorbance of eluents at 230 nm. The solutions collected from the HPLC runs were concentrated and lyophilized. Finally, purified conjugates were dissolved in sterile PBS (10 mM, pH 7.4) at concentrations between 1-2.5 mM and stored at -20 °C.

### TEM imaging of PA nanostructures

For TEM imaging, PAs conjugated with ICG were dissolved in ultrapure water at a concentration of 200 μM. Then, PA solutions (5 μL) were placed on carbonfilm 200 copper mesh TEM grids, incubated for 5 min before blotting, and air dried. For negative staining of the samples, a 2% uranyl acetate solution was prepared in ultrapure water and filtered using a 0.1 μm syringe filter. 20 μL of filtered uranyl acetate solution was then placed on a Parafilm piece and TEM grids were floated on this droplet for 7 min. Finally, TEM grids were blotted to remove excess stain and dried at room temperature. TEM imaging was performed using a Tecnai microscope (FEI).

### Stability of PA nanostructures in plasma

ICG conjugated PAs were dissolved in PBS (10 mM, pH 7.4) at a concentration of 1 or 100 μM, and 90 μL aliquots of these solutions were added to a 96 well plate (4 wells for each PA). Then, 10 μL of pooled human plasma was added to each well, and fluorescence spectra of ICG were recorded using a microplate reader (TECAN Spark 20M) at different time points.

### FPLC experiments

FPLC experiments were performed using SA-E, SA-K, or No-SA conjugated with ICG at a final concentration of 250 μM in filtered PBS (10 mM, pH 7.4) with or without human plasma or purified LPs or albumin. For analysis of SA-E interaction with purified LPs, VLDL, LDL, HDL, and human serum albumin were diluted in filtered PBS to approximate concentrations found in 10% human plasma: 0.0025 mg/mL, 0.13 mg/mL, 0.05 mg/mL, and 5 mg/mL, respectively. SA-E was incubated with the purified LP/protein mixture for 30 min at 37°C prior to injection. For analysis of plasma interactions, PAs or No-SA were incubated in PBS containing 10% pooled human plasma for 30 min at 37 °C prior to injection. The total volume injected for all samples was 400 μL. All data was collected on an KTA pure™ chromatography system using the following running conditions: size exclusion chromatography on a Superose 6 Increase 10/300 GL column with a column volume (CV) of 30 mL, pre-column pressure limit of 5 MPa, delta-column pressure limit of 2.6 MPa, flow rate of 0.75 mL/min, in 100% PBS. The sample running method included first an equilibration step prior to injection of the sample with 100% PBS for 0.5 CV, followed by sample application via 1 mL capillary loop in 100% PBS, and finished with a linear elution consisting of 2 CV collected in 1 mL fractions. The UV-Vis detector wavelengths used to detect eluting proteins were 280 nm and 700 nm to detect all proteins/LPs and ICG conjugated PAs, respectively.

### Mass Spectrometry

Samples were prepared for mass spectrometry analysis via a pull-down of a biotinylated version of the SA-E probe (SA-Eb, 100 μM) incubated with 50% pooled human plasma at 37 °C for 2 hours. Prior to incubation with the SA-Eb-plasma mixture, magnetic streptavidin-coated beads (Dynabeads MyOne Streptavidin T1, Invitrogen, 65601) were washed with PBS (10 mM, pH 7.4) then blocked with 1% casein equal to the total initial volume of beads for 15 min at room temperature. After blocking, beads were washed again with PBS. Beads were then combined with SA-Eb-plasma solution, 25 μL beads to 200 μL mixture. This mixture was then incubated at room temperature for 30 min. After incubation, the remaining solution was removed, and beads were washed five times with PBS. Bound SA-Eb was eluted from beads in 30 μL 0.5% Triton X-100 solution. Magnetic beads were then removed, and elution was collected for analysis. For plasma control, the incubation with SA-Eb step was skipped, and 50% plasma in PBS was incubated with beads and washed and eluted using the same method as the SA-Eb-plasma solution. The elution solutions were then analyzed by the Proteomics Shared Resource facility at Oregon Health & Science University (See Supporting Information for details).

### Cell lines

Cell lines used for the experiments were all tested for Mycoplasma at least once every other year. Following cell lines were used: HCT116 (human colorectal cancer, ATCC CCL-247), A375-Luc/iRFP (human melanoma, Creative Biogene CSC-RR0254), MCF7 (human breast cancer, ATCC HTB-22), 4T1 (mouse breast cancer, ATCC CRL-2539), RG2 (rat glioma, ATCC CRL-2433), and LL/2 (mouse lung cancer (LLC1), ATCC CRL-1642). MC38 mouse intestinal epithelial cancer cells were provided by Dr. Melissa Wong at OHSU. SR-B1-/- and wild type TRAMP-C2 mouse prostate cancer cells were provided by Jonathan D. Smith at Case Western Reserve University. LDL-R-/- and wild type HeLa human cervical cancer cells were purchased from Ubigene (YKO-H1027). Patient derived pancreatic or colon cancer cells were obtained at OHSU with patient consent and cultured for at least 5 passages. Cell lines were maintained in either RPMI Medium 1640 (Gibco, 11875093) supplemented with 10% fetal bovine serum (FBS) (Cytiva, SH30396-03) and 5% penicillin-streptomycin (pen-strep) (Gibco, 15140163) or DMEM with 4.5 g/L D-glucose, L-glutamine, and 110 mg/L sodium pyruvate (Gibco, 11995065) and supplemented with 10% FBS and 5% pen-strep as the growth media in the incubator at 37°C under 5% CO_2_. For mCherry transfection of RG2 cells, lentivirus targeting mCherry to the cell membrane (Takara, 0026VCT, rLV.EF1.mCherry-Mem-9) was used according to manufacturer’s recommendation. Transfected cells were passaged at least 10 times before being used for in vivo experiments.

### Cellular uptake studies

To compare the uptake of SA-E and SA-K, 1x10^5^ 4T1 cells were grown to near confluency (1-2 days) in 96 well plates with black walls and flat glass bottom. Cy5 conjugated PAs (20 µM) were preincubated in 10% FBS containing RPMI for 30 min at 37°C, wells were aspirated, and 100 µL of media containing PAs were added. Plate was incubated for 30 min at room temperature. Media was aspirated, wells were washed 3x with 150 µL sterile PBS and then 100 µL RPMI containing 10% FBS was added to each well. Plate was incubated for 1 hour at 37°C. Cell nuclei were stained with Hoechst (1:5000 dilution in PBS) for 5 min and then removed. 100 µL of phenol red free RPMI containing 10% FBS was added to each well. Confocal images were obtained with a Nikon CrestOptics X-Light V3 Spinning Disk Confocal microscope under 20x and 60x objective using NIS-Elements software (405 nm laser for Hoechst and 637 nm laser for Cy5 imaging). The microscope incubator was preheated for 30 minutes to reach 35 °C prior to imaging.

To study the uptake of SA-E, SA-K, or No-SA over time, 4T1 cells were grown to near confluency in a 96 well plate as described above. Cy5 conjugated PAs or No-SA (20 µM) were preincubated in 10% FBS containing RPMI for 30 min at 37°C. Cell nuclei were stained with Hoechst stain as above. Wells were aspirated and 100 µL of Cy5 conjugated SA-K or SA-E (20 µM) were added to wells for approximately 45 seconds. Wells were washed with sterile PBS (3x 150µL) and Confocal images were taken at different time points (up to 2 hours) after PA or No-SA addition as described above using a 60x objective.

To study the endocytosis of Cy5 conjugated SA-E, 4T1 cells were grown to near confluency in a 96 well plate as described above. FITC-dextran (2 mg/mL) and Cy5 conjugated SA-E (20 µM) were preincubated in 10% FBS containing RPMI for 30 min at 37°C. Wells were aspirated and 100 µL of media containing FITC-dextran and Cy5 conjugated SA-E was added. The plate was incubated at 37 °C for 4 hours. Wells were washed with PBS, nuclei were stained with Hoechst, and Confocal imaging was performed as described above. 477 nm laser was used for FITC imaging.

To study the effect of serum biomolecules on cellular uptake of SA-E, 4T1 cells were grown to near confluence in a 96 well plate as described above. Cy5 conjugated SA-E (20 µM) was preincubated in RPMI with or without 10% FBS for 30 min. Wells were aspirated, and 100 µL of media containing Cy5 conjugated SA-E was added. Then, the plate was incubated with SA-E and washed with PBS as described above. 100 µL of cell media, either with or without FBS as appropriate, were added to each well and the plate was returned to the incubator for 1 hour. To stain cell nuclei, a 1:5000 dilution of Hoechst stain was made in serum free media and added to each well. The plate was incubated for 5 min and fluorescence images were taken using a Leica Thunder DMi8 fluorescent microscope using LAS X software.

For SR-B1 studies, an equal number of wells in a black walled, glass bottom 96-well plate were seeded with either wild type or SR-B1-/- TRAMP-C2 cells and grown to near confluency, as above. The Cy5 conjugated SA-E was diluted in 10% FBS containing DMEM and incubated at room temperature for 30 min. All media from the wells was aspirated. Four wells of each cell type received 100 µL aliquots with SA-E (20 µM). The plate was incubated for 30 min, as above. Following the methods above, wells were washed with PBS, media was added, and the plate was returned to the incubator for 1 hour. Finally, Hoechst staining and fluorescence imaging were performed using a Leica Thunder DMi8 fluorescent microscope, as described above.

For LDL-R studies, 2x10^5^ wild type or LDL-R-/- HeLa human cervical cancer cells were grown to near confluency in 96 well plates with black walls and flat glass bottom. Cy5 conjugated SA-E (20 µM) was preincubated in DMEM containing 10% FBS for 30 minutes at 37°C. Wells were aspirated, and 100 µL of the SA-E containing DMEM was added. The plate was incubated for 30 min, as above. The wells were washed with sterile PBS, as above, media was added, and the plate was returned to the incubator for 1 hour. Wells were aspirated and nuclei were stained with Hoechst as above. Wells were aspirated and 100 µL of phenol red free DMEM with 10% FBS was added to each well. Cells were then imaged for Cy5 conjugated SA-E with a Zeiss Apotome 3 microscope at 20x and 63x objectives using Zen software.

To explore the effect of serum proteins and different inhibitors on the cellular uptake of SA-E, 4T1 cells were grown to near confluency in 96 well plates, as described above. For cellular uptake mechanism studies, the growth media (RPMI containing 10% FBS) was exchanged for 100 µL of FBS free media, then returned to the incubator. Inhibitors (brefeldin A, bromsulphalein, chlorpromazine hydrochloride, cis-ACBD, filipin, heparin, methyl-β-cyclodextrin, or wortmannin) were made in FBS free media at a concentration of 1 mM. Then, 1 µL of media containing inhibitors was added to wells, so that the final drug concentration in each well was 10 µM. The plate was returned to the incubator for 20 min. After incubation, Cy5 conjugated SA-E was added to all wells such that the final concentration was 20 µM per well. The plate was returned to the incubator for 30 min. Wells were aspirated and washed with PBS as above. 100 µL of FBS free media was added and the plate was returned to the incubator for 1 hour. Finally, Hoechst staining and fluorescence imaging were performed using a Leica Thunder DMi8 fluorescent microscope, as described above.

### Mouse tumor models

All experiments were approved by the Oregon Health and Science University (OHSU) Institutional Animal Care and Use Committee (IACUC) (TR01_IP00000674) and conformed to the guidelines set by the United States Animal Welfare Act and the National Institutes of Health. Mice were aged between 3-8 months and 20-30 g in weight and were housed in specific pathogen free cages. Human and rat cells were injected into either nude (The Jackson Laboratory, 002019) or NSG mice (The Jackson Laboratory, 005557). 4T1 cells were injected into Balb/c mice (The Jackson Laboratory, 000651). LLC1, MC38, and TRAMP-C2 were injected into C57Bl/6J mice (The Jackson Laboratory, 000664). Apc^min^ and MMTV-PyMT mice were purchased from the Jackson Laboratory (002020 and 002374). ApoA1 and LDL-R knockout mice were purchased from the Jackson Laboratory (002055 and 002207, respectively). Patient derived xenografts were generated by injecting the patient derived cancer cells subcutaneously in NSG mice.

### Blood circulation in mice

For blood circulation experiments, wild type mice were injected with 100 µL of ICG conjugated PAs or No-SA, or free ICG (0.5 mM). Then, blood was drawn via the saphenous vein or retro-orbitally at different time points (immediately after injection to 1 day). 2 µL of blood was diluted into 98 µL of PBS (10 mM, pH 7.4) and ICG fluorescence was measured using a microplate reader (TECAN Spark 20M) at excitation and emission wavelengths of 745 and 820 nm, respectively.

### Collection of blood components for FPLC and fluorescence measurements

For these experiments, 100 µL of ICG conjugated SA-E (0.5 mM) was intravenously injected into wild type (C57Bl/6J), ApoA1 -/-, or LDL-R -/- mice. One hour after injection blood was collected via cardiac puncture (500 µL) in EDTA containing tubes (BD MicroTainer Blood Collection Tube, 365974). To separate plasma, blood samples were centrifuged for 10 min at 1500 g, and supernatants were removed. Samples were centrifuged again, and supernatants were stored at -20°C for up to 5 days before use. For RBCs, 10 µL of the centrifugation pellet was separated and stored at -20 °C for up to 5 days before analysis. For WBCs, 100 µL of the remaining pellet was collected and combined with 2 mL Red Blood Cell Lysis Buffer (BioLegend, 420514). This mixture was allowed to incubate at room temperature for 15 min with occasional mixing. The solution was then centrifuged at 500 g for 5 min, the supernatant was removed, and the pellet was redissolved in 30 µL PBS, then stored at -20 °C for up to 5 days.

FPLC of plasma samples were performed as described above using an KTA pure™ chromatography system. For fluorescence measurements, 5 µL of samples were added to a 96 well plate, diluted to a total volume of 100 µL with PBS, and fluorescence was measured using a microplate reader (Tecan Spark 20M).

### Tumor accumulation in mice

For the following experiments, mice were injected intravenously with 100 µL of ICG conjugated PAs (0.5 mM) or free ICG (0.5 mM). For tumor accumulation studies, cells (0.5–1×10^6^ cells) were injected subcutaneously (HCT-116, A375, TRAMP-C2, MC38, LLC1, RG2, PDX cell lines, wild type or SR-B1 -/- TRAMP-C2, and wild type or LDL- R -/- HeLa cells), intradermally (A375 cells), or into the fat pad (4T1 and MCF-7 cells). Cells were then allowed to grow into tumor mass of ∼0.3-1 cm, typically 1-3 weeks. After probe injections, mice were imaged using the IVIS Spectrum (PerkinElmer) with excitation and emission wavelengths of 745 and 820 nm, respectively, at the specified time points. For MMTV mice experiments, mice were injected intravenously weekly with SA-E (100 μL, 0.5 mM) between 71 and 85 days of age and imaged 2 days later in the IVIS. For Apc^min^ mice experiments, mice were aged to 4 months to allow intestinal tumors to form. SA-E was injected intravenously (100 μL, 0.5 mM), 2 days later the intestinal tissue was harvested, washed and analyzed in the IVIS for ICG. For the ApoA1 and LDL-R experiments, wild type, ApoA1-/-, or LDL-R-/- mice were subcutaneously injected with syngeneic MC38 cancer cells. Tumors were allowed to grow to 0.5-1 cm and then injected intravenously with SA-E (100 μL, 0.5 mM). 2 days later, the mice were euthanized, and tumors and major organs were excised and ICG fluorescence measured using an IVIS (PerkinElmer). Living Image (PerkinElmer) software was used for all analyses.

### PA fluorescence in liver homogenates

Liver samples (6.2 g total weight) collected from healthy BALB/cJ mice were placed in a gentleMACS™ M Tube containing 2 mL of PBS. The M Tube was fitted with a 600 μm mesh strainer, and homogenization was performed using a gentleMACS Dissociator by running a one-minute program at 1260 rotations per round (rpr) for five cycles. ICG conjugated SA-E and SA-K were then spiked into the liver homogenates in a 96-well plate at final concentrations of 0, 0.1, 0.5, 2.0, 5.0, and 10 μM. Fluorescence signals from the homogenates were measured using an IVIS (PerkinElmer).

### Biodistribution in mice

For biodistribution studies, mice containing 4T1 tumors were injected intravenously with 100 µL of ICG conjugated PAs or free ICG (0.5 mM). 2 days after injection, the mice were euthanized, and tumors and organs were excised and analyzed in the IVIS (PerkinElmer) for ICG.

To determine the percentage of the injected dose of SA-E in tissues, tumor, spleen, kidney, lung, and colon tissues were harvested from 5 mice 2 days after ICG conjugated SA-E (IV, 100 µL, 0.5 mM). Organ weights were measured for normalization. Tissues were homogenized in gentleMACS™ M Tube containing 1 mL of PBS and equipped with a 600 μm mesh strainer using a gentleMACS dissociator as described above. The resulting tissue homogenates were transferred into a 96-well plate for fluorescence measurement using a microplate reader (TECAN Spark 20M). Additionally, organ tissues from healthy mice without probe injection were harvested and homogenized to prepare calibration curves. ICG conjugated SA-E was spiked into tissue samples at different concentrations, and fluorescence intensities were measured using a microplate reader (TECAN Spark 20M).

### Clearence study

Six Balb/c mice were intravenously injected with 100 µL of either ICG-conjugated SA-E or PBS, with three mice in each group. Urine and stool samples were collected from each mouse at different time points up to 7 days. Urine samples were diluted 10-fold in PBS, and ICG fluorescence was measured using a microplate reader (TECAN Spark 20M). Urine from healthy Balb/c mice was collected and pooled to generate a calibration curve. Fluorescence of stool samples were detected using an IVIS (PerkinElmer).

### Toxicity in mice

Wild type mice were injected intravenously with ICG conjugated SA-E at 4 times the dose used previously for imaging (100 µL, 2 mM). 1 day later, blood was removed via cardiac puncture, placed into an EDTA tube and then centrifuged at 1500g for 10 min, then 2500g for 10 min to obtain plasma. The plasma was analyzed for Aspartate Aminotransferase activity (AST) (Abcam, AB105135) and creatinine (Crystal Chem, 80350) following the manufacturer’s protocol. For complete blood counts, whole blood (10 µL) was mixed with 2 mM EDTA (10 µL) and analyzed using the HemaVet 950S (Drew Scientific).

### Flow cytometry

RG2-mCherry cells were implanted subcutaneously in nude mice and allowed to grow to 0.5-1 cm. Mice were then injected intravenously with the Cy5 labeled SA-E (100 µL, 0.5 mM). 2 days later, tumors were extracted, minced, and dissociated using 1 mg/mL solution collagenase/dispase (Roche, 10269638001 and 11097113001) for 45 min at 37 °C. The solution was broken down with pipetting and filtered through a 70 µm filter. The single-cell solution was suspended in sterile PBS (10 mM, pH 7.4) and then incubated with 1:50 dilution of AlexaFluor 488 Rat Anti-Mouse CD45 antibody (BD Pharminogen, 567377) for 45 min at room temperature. Finally, the cells were pelleted and, washed and incubated with 5 µM Calcein Blue, AM (Invitrogen, C1429) for 20 min. The cells were then analyzed in BD FACSymphony in the DAPI channel (355nm laser, 450/50 filter), FITC channel (488 nm laser, 515/30 filter), Cy3 channel (561 nm laser, 586/15 filter), and Cy5 channel (628 nm laser, 670/30 filter). Cells in culture were used to establish size and mCherry intensity. Tumors from mice that did not receive any SA-E injection and unstained cells were used as controls for gating of SA-E and CD45. Finally, three experimental mice were analyzed in all channels using the gating established from the controls. Gating was performed in FlowJo software.

### Rat models

The care and use of animals were approved by the Institutional Animal Care and Use Committee and were supervised by the OHSU Department of Comparative Medicine (DCM) (protocol IP00000843). Female Long Evans rats (200-300g) were purchased from Charles River Laboratories by OHSU DCM. Rats were anesthetized using 3% isoflurane and a 2-mm diameter hole was drilled in the skull using a 27-gauge needle in a stereotactic frame (David Kopf Instruments; Tujunga, CA). Cancer cells (10^6^ cells/10 μL, >90% viability) were injected at a rate of 1 μL/min. Rat RG2 cells with or without mCherry were implanted into the right caudate nucleus (bregma= 0; lateral 0.31 cm; vertical - 0.65 cm).

### MRI of rat tumors

MRI was performed to confirm tumor growth prior to administration of SA-E using a horizontal bore 11.75 T magnet (Bruker Scientific Instruments, Billerica MA) maintained by OHSU Advanced Imaging Research Center (AIRC). Animals were imaged 8-35 days after tumor implantation, depending on tumor type. Under intraperitoneal dexmedetomidine (Zoetis, 0.5 mg/kg) and ketamine (Zoetis, 60 mg/kg) sedation, the lateral caudal vein was catheterized and the animal was securely placed in a Plexiglas holder and positioned in the small animal MRI. T1 (Fat saturated 2D Spine Echo [SE], TR 160, TE 1.654), T2 (2D SE, TR 4020.62, TE 23.64) and T2* (2D Gradient Echo, TR 430.25, TE 6.656, FA 30) sequences were obtained prior to a dose of 50 μmol/kg of Omniscan via tail vein catheter followed by a ∼0.25 mL saline flush. A post-contrast T1 scan was repeated 5 minutes following the administration of Omniscan.

### IVIS and Confocal imaging of rat tumors

After confirmation of RG2 tumor growth, ICG conjugated SA-E (1 mL, 0.5 mM) was delivered via tail vein catheter, followed by a ∼0.25 mL saline flush, prior to anesthetic reversal with intraperitoneal atipamezole HCl (Antisedan, 1.0 mg/kg). For IVIS imaging of rat tumors, animals were euthanized with euthasol (Virbac) via intracardiac injection under isoflurane anesthetization 2 days after SA-E administration. Brain, heart, lung, liver, spleen and kidney were harvested and stored in PBS until IVIS imaging (∼30 min after resection). Tissues were first imaged in the IVIS for ICG signal. Then, wholemount brain sections were imaged under IVIS for the ICG signal of SA-E and the mCherry signal of RG2 cells. For Confocal imaging of brain sections, the brain was bisected through the tumor, flash frozen and stored at -80 °C in O.C.T. embedding medium (Sakura Finetek USA Inc, 4583). The tissue was then sectioned with a cryostat (Leica CM 3050s) at 15 μm thickness and stored on dried ice prior to fixing in 4% formaldehyde for 5 min. Tissue sections were rinsed in PBS and then counterstained with Hoechst stain (1:4000 dilution in PBS) for 5 min and imaged on a Nikon CrestOptics X-Light V3 Spinning Disk Confocal microscope (Hoechst 405 nm laser, mCherry 545 nm laser, ICG 748 nm laser) with 20x and 60x objective using NIS-Elements software.

### Cytotoxicity of SA-E-MMAE

To assess the cytotoxicity of SA-E-MMAE and free MMAE, 4T1 cells, and RG2 cells were seeded in 96-well plates at a density of 5×10^3^ cells per well in 100 μL of RPMI (10% FBS). The plates were then incubated at 37 °C under 5% CO_2_ for 1 day prior to the addition of SA-E-MMAE or free MMAE, resulting in final drug concentrations ranging from 0 to 10 μM for SA-E-MMAE and 0 to 0.05 μM for MMAE. Plates were further incubated at 37 °C under 5% CO_2_ for 3 days. To evaluate cellular viabilities, the RPMI media in each well was replaced with fresh RPMI containing 10% MTS reagent. Subsequently, plates were incubated at 37 °C for 2 hours before measuring the MTS absorbance at 490 nm using a microplate reader (TECAN Spark 20M).

### Antitumor effect of SA-E-MMAE

Mice were given 5×10^5^ cells injected into either the fat pad (4T1 in Balb/c mice) or subcutaneously (RG2 or HCT-116 in nude mice). 4T1 cells and RG2 cells were injected in PBS and HCT-116 were injected in 1:1 mixture of PBS and Matrigel (Life Science 354263). Mice were then treated intravenously with the free MMAE (0.1 or 0.3 mg/kg), SA-E-MMAE (0.3, 0.6, 0.8, or 1 mg/kg) or PBS (10 mM, pH 7.4). 5 doses were given every 2-3 days over 11 days. For 0.3 mg/kg dose of free MMAE, the drug was administered 3 times as a significant weight loss was observed after the third injection. Mouse weight was monitored throughout the study. Tumor size was measured in two directions with a caliper 2-3 times per week rounded to the nearest tenth of a mm. Volume was calculated as a basic sphere of 4/3πr^3^.

## Data availability statement

All the data of this study are available within the paper, extended figures, Supporting information, or upon request from the corresponding authors.

## Acknowledgements

Authors thank Drs. Melissa Wong and Jonathan D. Smith for providing MC38 and TRAMP-C2 (wild type and SR-B1 -/-) cells, respectively. Authors thank Randall J. Armstrong for his help with FPLC measurements. Mass spectrometric analysis was performed by the OHSU Proteomics Shared Resource with partial support from NIH core grants P30EY010572, P30CA069533 and OHSU Emerging Technology Fund. Author Ramon F. Barajas Jr is supported by NIH NCI 1K08CA237809 and L30 CA220897. Leslie Muldoon and Ramon F Barajas Jr. are supported by Jonathan D + Mark C. Lewis Foundation. This project was supported by funding from the NIH (R21NS135452) and the Cancer Early Detection Advanced Research (CEDAR) center at the Oregon Health & Science University’s Knight Cancer Institute.

## Author contributions

A.Y. designed the PAs. J.M.F. and A.Y. designed the study. L.L.M. and R.F.B. designed rat studies. B.P.B. provided insights on the concept. L.X. and M.D. performed PA conjugations and purification. M.R.S. performed fluorescence and FPLC experiments. M.R.S. and F.G. performed sample preparation for MS measurements. S.S. performed TEM imaging. I.D. and S.R. designed and conducted MD simulations. K.M., J.M.F., S.D., B.R.K., and K.B. performed cellular uptake experiments. L.X., M.R.S., J.M.F., A.Q., S.D., and X.Y. performed mouse studies. K.M., J.M.F. and S.H. performed rat studies. L.X. performed cytotoxicity studies. J.M.F. and A.Y. wrote the manuscript with input from all authors.

## Competing interests

A.Y., J.M.F., and B.P.B submitted a patent application (PCT/US2022/073964) on the early findings of this study. The other authors declare no competing interests.

## Notes

### Summary of Updates

The revised manuscript includes more in vivo data on the mechanism of cancer-specific accumulation of the peptide amphiphiles developed in this study. In addition, more in vitro studies were performed to understand the cellular internalization of peptide amphiphiles.

## References

1. Levin, A.; Aida, T.; Hamley, S.; Castelletto, V.; Pashkina, E. D. S.; Palikov, V. A.; Blokhin, K. A.; Reza, M.; Ruokolainen, J.; Hamley, I. W. Biomimetic Peptide Self-Assembly for Functional Materials. Nat. Rev. Chem. 2020, 4, 615–634.

2. Vicente-García, C.; Colomer, I. Lipopeptides as Tools in Catalysis, Supramolecular, Materials and Medicinal Chemistry. Nat. Rev. Chem. 2023, 7, 710–731.

3. Zhao, X.; Pan, F.; Xu, H.; Yaseen, M.; Shan, H.; Hauser, C. A. E.; Zhang, S.; Lu, J. R. Molecular Self-Assembly and Applications of Designer Peptide Amphiphiles. Chem. Soc. Rev. 2010, 39, 3480–3498.

4. Nagarajan, R. Molecular Packing Parameter and Surfactant Self-Assembly: The Neglected Role of the Surfactant Tail. Langmuir 2002, 18, 31–38.

5. Zhang, S. Fabrication of Novel Biomaterials Through Molecular Self-Assembly. Nat. Biotechnol. 2003, 21, 1171–1178.

6. Sinha, N. J.; Langenstein, M. G.; Pochan, D. J.; Kloxin, C. J.; Saven, J. G. Peptide Design and Self-assembly into Targeted Nanostructure and Functional Materials. Chem. Rev. 2021, 121, 13915–13935.

7. Rad-Malekshahi, M.; Lempsink, L.; Amidi, M.; Hennink, W. E.; Mastrobattista, E. Biomedical Applications of Self-Assembling Peptides. Bioconjug. Chem. 2016, 27, 3–18.

8. Zhou, Y.; Chen, X.; Cao, J.; Gao, H.; Zhang, W.; Loh, X. J. Molecularly Stimuli-Responsive Self-Assembled Peptide Nanoparticles for Targeted Imaging and Therapy. ACS Nano 2023, 17, 8004–8025.

9. Eskandari, S.; Guerin, T.; Toth, I.; Stephenson, R. J. Recent Advances in Self-Assembled Peptides: Implications for Targeted Drug Delivery and Vaccine Engineering. Adv. Drug Deliv. Rev. 2017, 110–111, 169–187.

10. Acar, H.; Ting, J. M.; Srivastava, S.; LaBelle, J. L.; Tirrell, M. V. Self-assembling Peptide-based Building Blocks in Medical Applications. Adv. Drug Deliv. Rev. 2017, 110–111, 65–79.

11. Chung, E. J.; Cheng, Y.; Morshed, R.; Nord, K.; Han, Y.; Wegscheid, M. L.; Auffinger, B.; Wainwright, D. A.; Lesniak, M. S. Fibrin-binding, Peptide Amphiphile Micelles for Targeting Glioblastoma. Biomaterials 2014, 35, 1249–1256.

12. Trent, A.; Marullo, R.; Lin, B.; Black, M.; Tirrell, M. Structural Properties of Soluble Peptide Amphiphile Micelles. Soft Matter 2011, 7, 9572–9582.

13. Pitman, M.; Larsen, J. The Characterization of Self-Assembled Nanostructures in Whole Blood. Anal. Methods 2020, 12, 2068–2081.

14. Dube, N.; Seo, J. W.; Dong, H.; Shu, J. Y.; Lund, R.; Mahakian, L. M.; Ferrara, K. W. Effect of Alkyl Length of Peptide–Polymer Amphiphile on Cargo Encapsulation Stability and Pharmacokinetics of 3-Helix Micelles. Biomacromolecules 2014, 15, 2963–2970.

15. Tsonchev, S.; Troisi, A.; Schatz, G. C.; Ratner, M. A. On the Structure and Stability of Self-Assembled Zwitterionic Peptide Amphiphiles: A Theoretical Study. Nano Lett. 2004, 4, 427– 431.

16. Black, M.; Trent, A.; Kostenko, Y.; Lee, J. S.; Olive, C.; Tirrell, M. Self-Assembled Peptide Amphiphile Micelles Containing a Cytotoxic T-Cell Epitope Promote a Protective Immune Response In Vivo. Adv. Mater. 2012, 24, 3845–3849.

17. Monopoli, M. P.; Åberg, C.; Salvati, A.; Dawson, K. A. Biomolecular Coronas Provide the Biological Identity of Nanosized Materials. Nat. Nanotechnol. 2012, 7, 779–786.

18. Dawson, K. A.; Yan, Y. Current Understanding of Biological Identity at the Nanoscale and Future Prospects. Nat. Nanotechnol. 2021, 16, 229–242.

19. Mahmoudi, M.; Landry, M. P.; Moore, A.; Coreas, R. The Protein Corona from Nanomedicine to Environmental Science. Nat. Rev. Mater. 2023, 8, 422–438.

20. Busatto, S.; Pham, A.; Suh, J.; Shapiro, S. Lipoprotein-based Drug Delivery. Adv. Drug Deliv. Rev. 2020, 159, 377–390.

21. Thaxton, C. S.; Rink, J. S.; Naha, P. C.; Cormode, D. P. Lipoproteins and lipoprotein mimetics for imaging and drug delivery. Adv. Drug Deliv. Rev. 2016, 106, 116–131.

22. Mulder, W. J. M.; Jaffer, F. A.; Fayad, Z. A.; Nahrendorf, M. High-Density Lipoprotein Nanobiologics for Precision Medicine. Acc. Chem. Res. 2018, 51, 127–137.

23. Kuai, R.; Li, D.; Chen, Y. E.; Moon, J. J.; Schwendeman, A. High-Density Lipoproteins: Nature’s Multifunctional Nanoparticles. ACS Nano 2016, 10, 3015–3041.

24. Paramonov, S. E.; Jun, H.-W.; Hartgerink, J. D. Self-Assembly of Peptide−Amphiphile Nanofibers: The Roles of Hydrogen Bonding and Amphiphilic Packing. J. Am. Chem. Soc. 2006, 128, 7291–7298.

25. Koda, D.; Maruyama, T.; Minakuchi, N.; Nakashima, K.; Goto, M. Proteinase-mediated drastic morphological change of peptide-amphiphile to induce supramolecular hydrogelation. Chem. Commun. 2010, 46, 979–981.

26. Versluis, F.; Marsden, H. R.; Kros, A. Power struggles in peptide-amphiphile nanostructures. Chem. Soc. Rev. 2010, 39, 3434–3444.

27. Mundra, V.; Peng, Y.; Rana, S.; Natarajan, A.; Mahato, R. I. Micellar formulation of indocyanine green for phototherapy of melanoma. J. Control. Release 2015, 220, 130–140.

28. Missirlis, D.; Krogstad, D. V.; Tirrell, M. Internalization of p5314-29 peptide amphiphiles and subsequent endosomal disruption results in SJSA-1 cell death. Mol. Pharm. 2010, 7, 2173–2184.

29. Lu, J.; Owen, S. C.; Shoichet, M. S. Stability of Self-Assembled Polymeric Micelles in Serum. Macromolecules 2011, 44, 6002–6008.

30. Kratz, F. A clinical update of using albumin as a drug vehicle - A commentary. J. Control. Release 2014, 190, 331–336.

31. Zorzi, A.; Middendorp, S. J.; Wilbs, J.; Deyle, K.; Heinis, C. Acylated heptapeptide binds albumin with high affinity and application as tag furnishes long-acting peptides. Nat. Commun. 2017, 8, 16092.

32. Sarett, S. M.; Werfel, T. A.; Lee, L.; Jackson, M. A.; Kilchrist, K. V.; Brantley-Sieders, D. M.; Duvall, C. L. Lipophilic siRNA targets albumin in situ and promotes bioavailability, tumor penetration, and carrier-free gene silencing. Proc. Natl. Acad. Sci. 2017, 114, E6490–E6497.

33. Zhu, G.; Lynn, G. M.; Jacobson, O.; Chen, K.; Liu, Y.; Zhang, H.; Ma, Y.; Zhang, F.; Tian, R.; Ni, Q.; Cheng, S.; Wang, Z.; Lu, N.; Yung, B. C.; Wang, Z.; Lang, L.; Fu, X.; Jin, A.; Weiss, I. D.; Vishwasrao, H.; Niu, G.; Shroff, H.; Klinman, D. M.; Seder, R. A.; Chen, X. Albumin/vaccine nanocomplexes that assemble in vivo for combination cancer immunotherapy. Nat. Commun. 2017, 8, 1954.

34. Ren, L.; Yi, J.; Yang, Y.; Li, W.; Zheng, X.; Liu, J.; Li, M.; Wang, Z.; Liu, H.; Li, J.; Cheng, S.; Zhang, J.; Xu, Z. Apolipoproteins and cancer. Cancer Med. 2019, 8, 7032–7043.

35. Blair, H. C.; Kalyvioti, E.; Papachristou, N. I.; Tournis, S.; Syggelos, S. A.; Deligianni, D.; Orhii, P. B.; Zaidi, M. Apolipoprotein A-1 regulates osteoblast and lipoblast precursor cells in mice. Lab. Invest. 2016, 96, 763–772.

36. Evangelho, J. S.; Gomes, G. N.; Pereira, A. C.; Mill, J. G.; Baldo, M. P. Hypercholesterolemia magnitude increases sympathetic modulation and coagulation in LDLr knockout mice. Auton. Neurosci. 2011, 159, 98–103.

37. Yamashita, A.; Singh, S. P.; Hossain, M. A.; Choudhury, M. G. R.; Wakabayashi, H.; Tomita, Y.; Takikawa, M.; Iijima, H.; Yoshimoto, S.; Yoshiya, S.; Shimizu, M.; Tanaka, M.; Kuniyil, A.; Shibuya, S.; Soga, T.; Watanabe, M. Indocyanine Blue (ICB) as a Functional Alternative to Indocyanine Green (ICG) for Enhanced 700 nm NIR Imaging. Int. J. Mol. Sci. 2024, 25, 13547.

38. Karagiannis, E. D.; Urbanska, A. M.; Sahay, G.; Pelet, J. M.; Jhunjhunwala, S.; Langer, R.; Anderson, D. G. Rational Design of a Biomimetic Cell Penetrating Peptide Library. ACS Nano 2013, 7, 8616–8626.

39. Missirlis, D.; Khant, H.; Tirrell, M. Mechanisms of Peptide Amphiphile Internalization by SJSA-1 Cells in Vitro. Biochemistry 2009, 48, 3304–3314.

40. Missirlis, D.; Teesalu, T.; Black, M.; Tirrell, M. The Non-Peptidic Part Determines the Internalization Mechanism and Intracellular Trafficking of Peptide Amphiphiles. PLOS ONE 2013, 8, e54611.

41. Missirlis, D.; Krogstad, D. V.; Tirrell, M. Internalization of p5314-29 Peptide Amphiphiles and Subsequent Endosomal Disruption Results in SJSA-1 Cell Death. Mol. Pharm. 2010, 7, 2173–2184.

42. Kozarsky, K. F.; Donahee, M. H.; Rigotti, A.; Iqbal, S. N.; Edelman, E. R.; Krieger, M. Overexpression of the HDL receptor SR-BI alters plasma HDL and bile cholesterol levels. Nature 1997, 387, 414–417.

43. Traughber, C. A.; Opoku, E.; Brubaker, G.; Major, J.; Lu, H.; Lorkowski, S. W.; Gulshan, K. Uptake of high-density lipoprotein by scavenger receptor class B type 1 is associated with prostate cancer proliferation and tumor progression in mice. J. Biol. Chem. 2020, 295, 8252–8261.

44. Conner, S. D.; Schmid, S. L. Regulated portals of entry into the cell. Nature 2003, 422, 37–44.

45. Banushi, B.; Joseph, S. R.; Lum, B.; Lee, J. J.; Simpson, F. Endocytosis in cancer and cancer therapy. Nat. Rev. Cancer 2023, 23, 450–473.

46. Chen, S.; Zhong, Y.; Fan, W.; Xiang, J.; Wang, G.; Zhou, Q.; Wang, J.; Geng, Y.; Sun, R.; Zhang, Z.; Piao, Y.; Wang, J.; Zhuo, L.; Li, W.; Zheng, C.; Zhao, Y. Enhanced tumour penetration and prolonged circulation in blood of polyzwitterion-drug conjugates with cell-membrane affinity. *Nat*. Biomed. Eng. 2021, 5, 1019–1037.

47. Chakraborty, A.; Jana, N. R. Clathrin to Lipid Raft-Endocytosis via Controlled Surface Chemistry and Efficient Perinuclear Targeting of Nanoparticle. J. Phys. Chem. Lett. 2015, 6, 3688–3697.

48. Kanatani, I.; Ikai, T.; Okazaki, A.; Jo, J.; Yamamoto, M.; Imamura, M.; Kanematsu, A.; Yamamoto, Y.; Ito, N.; Ogawa, O.; Tabata, Y. Efficient gene transfer by pullulan-spermine occurs through both clathrin- and raft/caveolae-dependent mechanisms. J. Control. Release 2006, 116, 75–82.

49. Sezgin, E.; Levental, I.; Mayor, S.; Eggeling, C. The mystery of membrane organization: composition, regulation and roles of lipid rafts. Nat. Rev. Mol. Cell Biol. 2017, 18, 361–374.

50. Sviridov, D.; Mukhamedova, N.; Miller, Y. I. Lipid rafts as a therapeutic target: Thematic Review Series: Biology of Lipid Rafts. J. Lipid Res. 2020, 61, 687–695.

51. Li, B.; Qin, Y.; Yu, X.; Xu, X.; Yu, W. Lipid raft involvement in signal transduction in cancer cell survival, cell death and metastasis. Cell Prolif. 2022, 55, e13167.

52. Campiani, G.; Morelli, E.; Nacci, V.; Fattorusso, C.; Ramunno, A.; Novellino, E.; Greenwood, J.; Griffiths, R.; Sinclair, C.; McArthur, R. A.; di Renzo, G.; Mennini, T. A Rational Approach to the Design of Selective Substrates and Potent Nontransportable Inhibitors of the Excitatory Amino Acid Transporter EAAC1 (EAAT3). New Glutamate and Aspartate Analogues as Potential Neuroprotective Agents. J. Med. Chem. 2001, 44, 2507–2510.

53. Usama, S. M.; Zhao, B.; Burgess, K. Role of Albumin in Accumulation and Persistence of Tumor-Seeking Cyanine Dyes. Bioconjug. Chem. 2020, 31, 248–259.

54. Feingold, K. R. Lipid and Lipoprotein Metabolism. Endocrinol. Metab. Clin. 2022, 51, 437–458.

55. Attalla, S.; Taifour, T.; Bui, T.; Muller, W. Insights from transgenic mouse models of PyMT-induced breast cancer: recapitulating human breast cancer progression in vivo. Oncogene 2021, 40, 475–491.

56. Moser, A. R.; Pitot, H. C.; Dove, W. F. ApcMin: A mouse model for intestinal and mammary tumorigenesis. Eur. J. Cancer 1995, 31, 1061–1064.

57. Hingorani, D. V.; Doan, M. K.; Camargo, M. F.; Aguilera, J.; Song, S. M.; Pizzo, D.; Scanderbeg, D. J.; Cohen, E. E. W.; Lowy, A. M.; Adams, S. R. Monomethyl auristatin antibody and peptide drug conjugates for trimodal cancer chemo-radio-immunotherapy. Nat. Commun. 2022, 13, 3869.

