## Supporting Information for "Peptide Amphiphiles Hitchhike on Endogenous Biomolecules for Enhanced Cancer Imaging and Therapy"

**Supplementraty Data**

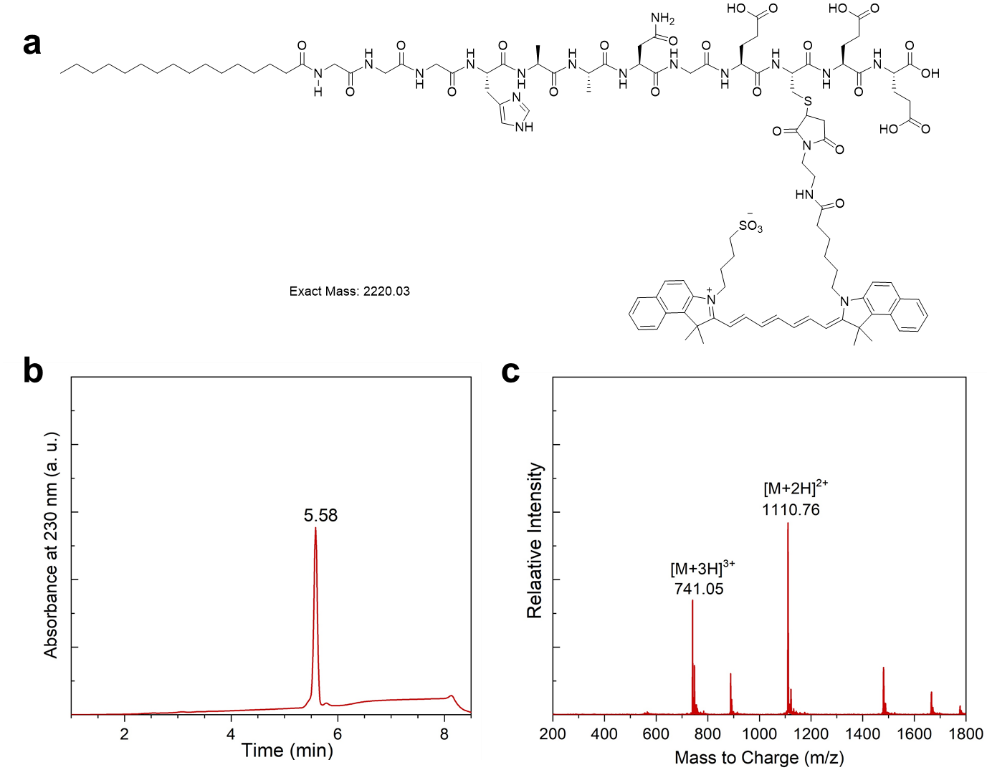

**Figure S1. Synthesis of ICG conjugated SA-E.** a) Molecular structure of ICG conjugated SA-E. b) LC and c) MS traces of the HPLC purified product.

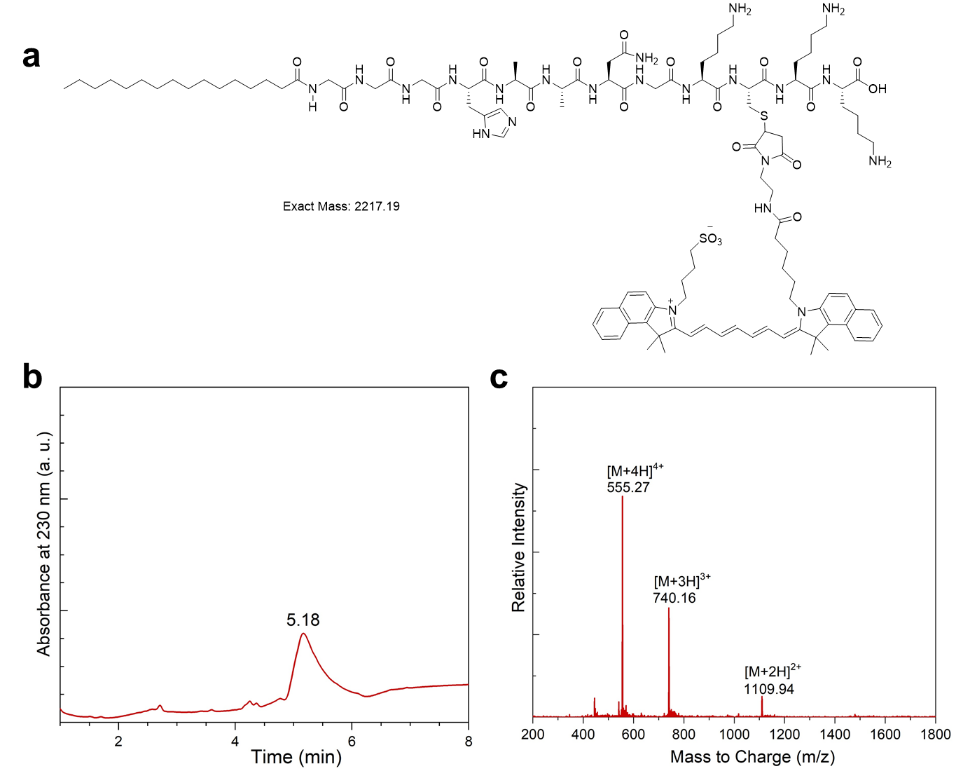

**Figure S2. Synthesis of ICG conjugated SA-K.** a) Molecular structure of ICG conjugated SA-K. b) LC and c) MS traces of the HPLC purified product.

**Molecular Dynamic (MD) Simulations:** For MD simulations, we modeled each PA into tube-like structures. Each tube-like structure was made by stacking 17 rings of 8 peptides each. These structures comprised of 264 peptides each to allow the formation of macromolecular structures. Then, MD simulations were performed, and the PA structures were observed over a ~0.83 µs simulation time. Though the structures have the same starting configuration (Figure 3a), we observed two distinct patterns at the end of simulations (Figure 3b). SA-E formed several spherical-like clumps while SA-K stayed in the tube formation, just compacted.

Figure 3c shows the root mean square deviation (RMSD) of the PA structures through the course of the simulations. While both SA-E and SA-K show typical equilibration behavior over the first 10-20 ns, SA-E showed a larger deviation from the starting tube-like orientation compared to the SA-K. This is in accordance with the snap-shots obtained at ~830 ns, where SA-K assembled into tube-like structures, and SA-E broke apart into smaller clumps and appeared to form micellar structures (Figure 3b). Figure 3d shows the number of contacts (normalized to per PA) between the heavy atoms of PAs. Consistent with the tube-like structures, which lead to a larger macromolecular assembly, we found a distinctly high number of interatomic contacts for the SA-K compared to the SA-E. In summary, SA-E groups together have both higher RMSD and a lower number of contacts. This is due to the spherical clumping behavior. When in spherical clumps, the PAs have moved farther away from their original cylindrical shape and are making fewer contacts due to there being a lower number of peptides in each sphere. The inverse occurred for SA-K due to the PAs staying in a cylindrical tube-like shape with more contacts. Overall, these results suggested that SA-K formed more strongly assembled structures than SA-E.

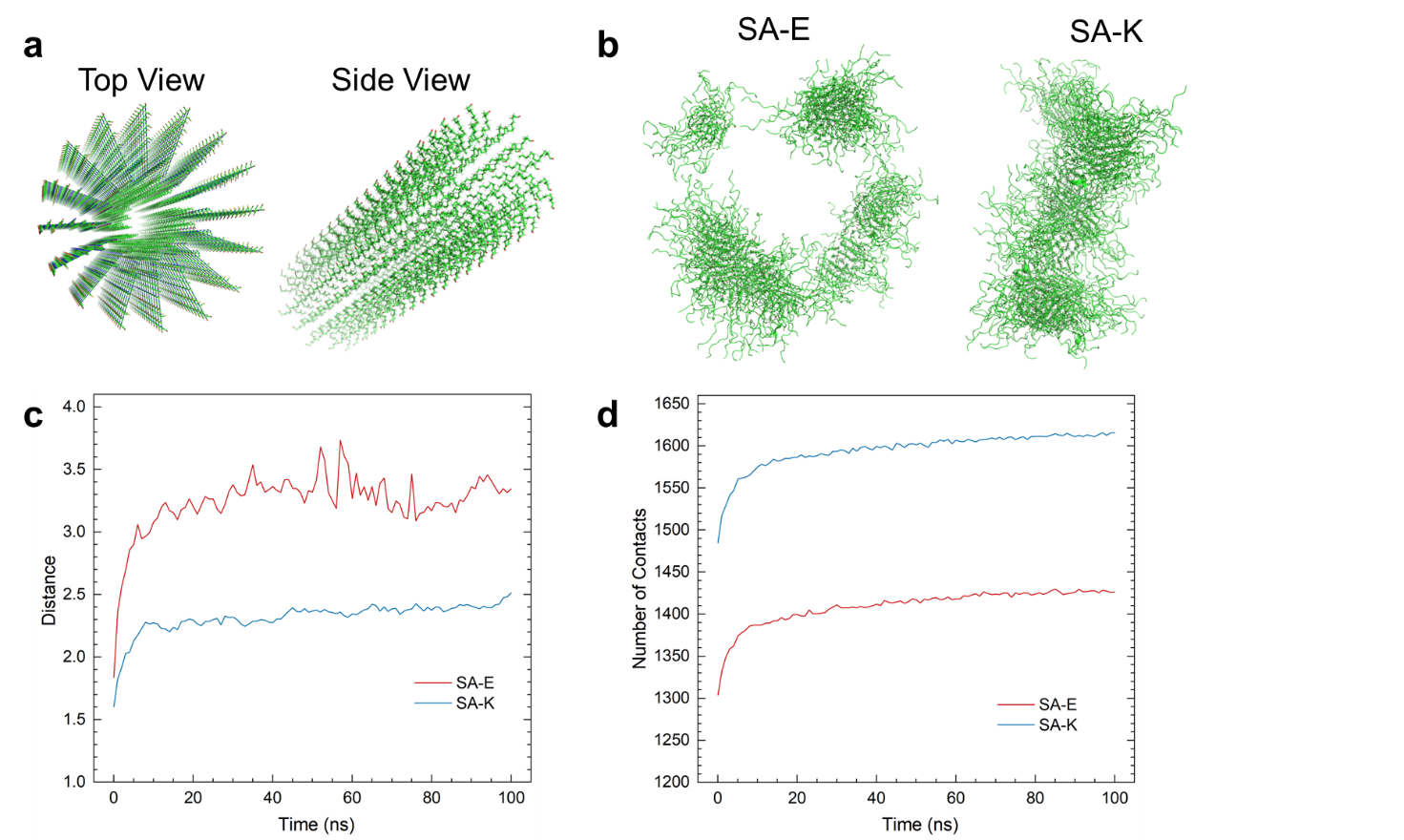

**Figure S3. Molecular dynamics simulations.** a) Representative images showing the initial PA structure used in simulations. b) Snap-shot images showing the final structures for each PA. c) RMSD distance (nm) and d) number of contacts of each PA over the course of simulations.

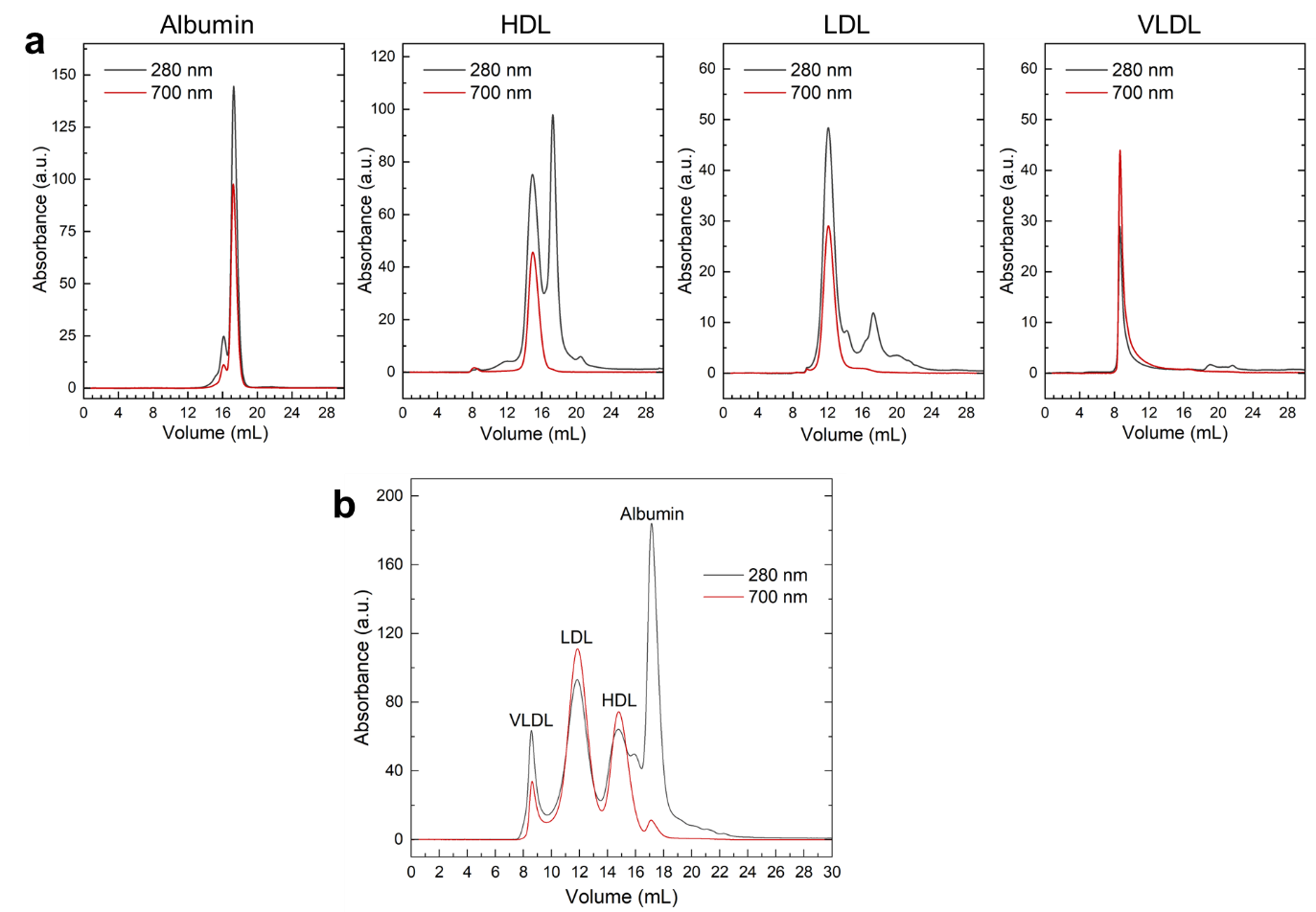

**Figure S4. SA-E assembles with albumin and LPs.** FPLC trace of SA-E in PBS containing human serum albumin (HSA), HDL, LDL, or VLDL (a) or a mixture of HSA and LPs. Absorbance at 280 nm was used to detect LPs and albumin, and 700 nm was used to detect SA-E. Albumin impurity was detected in LPs, especially for HDL. SA-E was not present in the albumin impurity peaks, further suggesting the stronger affinity of SA-E against LPs than albumin.

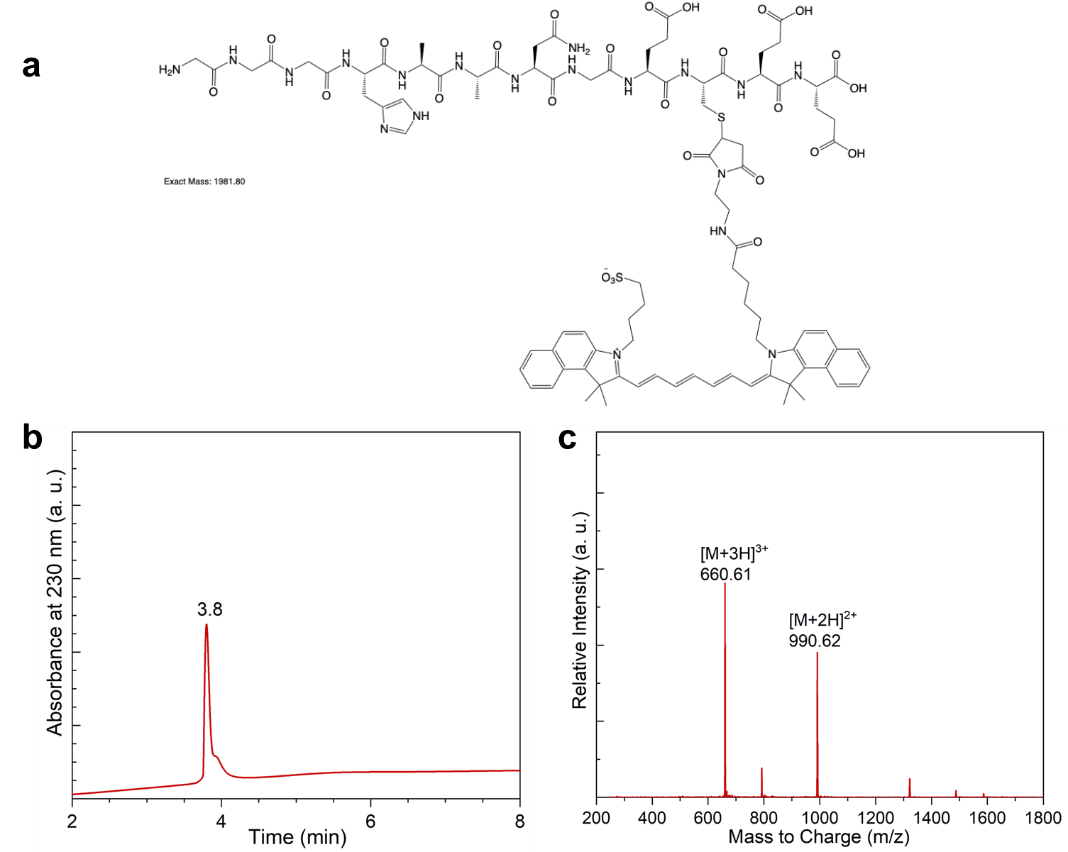

**Figure S5. Synthesis of ICG conjugated No-SA.** a) Molecular structure of ICG conjugated SA-Eb. b) LC and c) MS traces of the HPLC purified product.

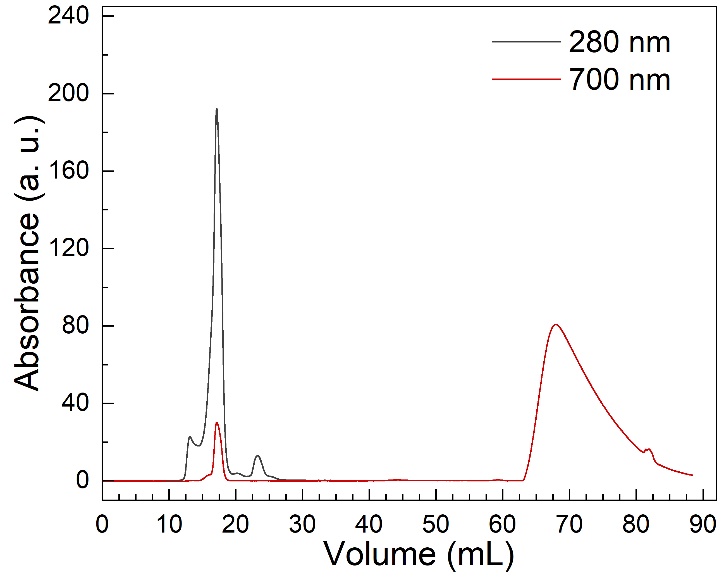

**Figure S6. No-SA did not bind to plasma components.** FPLC traces of ICG labeled No-SA in 10% human plasma.

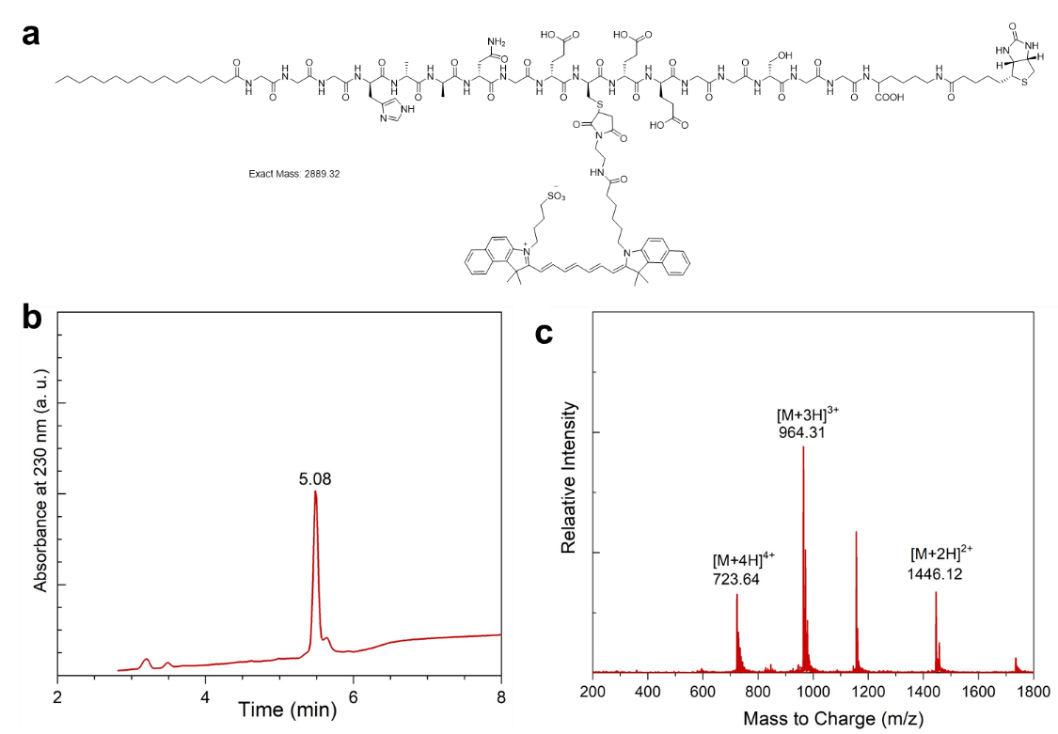

**Figure S7. Synthesis of ICG conjugated SA-Eb.** a) Molecular structure of ICG conjugated SA-Eb. b) LC and c) MS traces of the HPLC purified product.

**Supporting Table 1.** Summary of the proteins detected using MS in the presence or absence of biotinylated SA-E in human plasma.

|  | **Total Counts** | | | |  | **Percentage (%)** | | | |
| --- | --- | --- | --- | --- | --- | --- | --- | --- | --- |
| **Proteins** | **SA-E** | **SA-E** | **Only plasma** | **Only plasma** |  | **SA-E** | **SA-E** | **Only plasma** | **Only plasma** |
| **Albumin** | 1215 | 923 | 486 | 343 |  | 34.0 | 32.5 | 32.6 | 31.0 |
| **Apolipoproteins** | 891 | 909 | 171 | 134 |  | 25.0 | 32.0 | 11.5 | 12.1 |
| *Apolipoprotein A-I* | *560* | *600* | *109* | *80* |  | *15.7* | *21.1* | *7.3* | *7.2* |
| *Apolipoprotein B-100* | *166* | *170* | *0* | *0* |  | *4.7* | *6.0* | *0.0* | *0.0* |
| *Apolipoprotein E* | *66* | *59* | *25* | *24* |  | *1.8* | *2.1* | *1.7* | *2.2* |
| *Apolipoprotein A-IV* | *39* | *16* | *19* | *18* |  | *1.1* | *0.6* | *1.3* | *1.6* |
| *Apolipoprotein D* | *22* | *22* | *8* | *5* |  | *0.6* | *0.8* | *0.5* | *0.5* |
| *Apolipoprotein A-II* | *6* | *6* | *1* | *0* |  | *0.2* | *0.2* | *0.1* | *0.0* |
| *Apolipoprotein C-III* | *4* | *5* | *6* | *5* |  | *0.1* | *0.2* | *0.4* | *0.5* |
| *Apolipoprotein M* | *11* | *12* | *1* | *0* |  | *0.3* | *0.4* | *0.1* | *0.0* |
| *Apolipoprotein C-I* | *6* | *9* | *2* | *2* |  | *0.2* | *0.3* | *0.1* | *0.2* |
| *Apolipoprotein L1* | *7* | *7* | *0* | *0* |  | *0.2* | *0.2* | *0.0* | *0.0* |
| *Apolipoprotein(a)* | *4* | *3* | *0* | *0* |  | *0.1* | *0.1* | *0.0* | *0.0* |
| **Immunoglobulins** | 444 | 421 | 272 | 236 |  | 12.4 | 14.8 | 18.3 | 21.3 |
| **Other** | 1019 | 588 | 560 | 393 |  | 28.6 | 20.7 | 37.6 | 35.5 |
| **Total** | 3569 | 2841 | 1489 | 1106 |  | 100 | 100 | 100 | 100 |

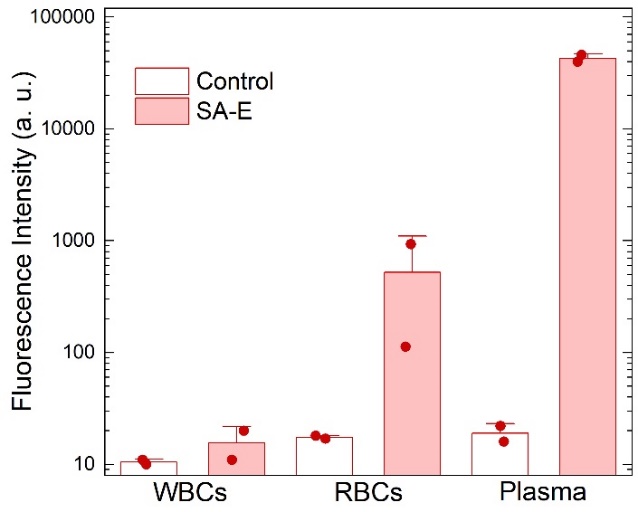

**Figure S8. SA-E mainly assembles with plasma biomolecules.** Total ICG labeled SA-E fluorescence in the blood plasma, WBC, or RBC components of mouse blood collected 1 hour after SA-E (50 nmole) injection. Data are presented as mean ± SEM.

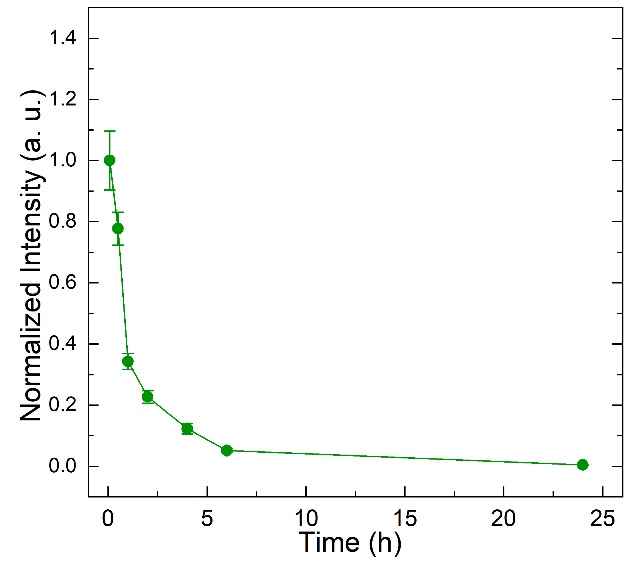

**Figure S9. Blood circulation of No-SA** in wild type mice showing that lipid modification (SA-E, Figure 1i) improves blood circulation. Data are presented as mean ± SEM.

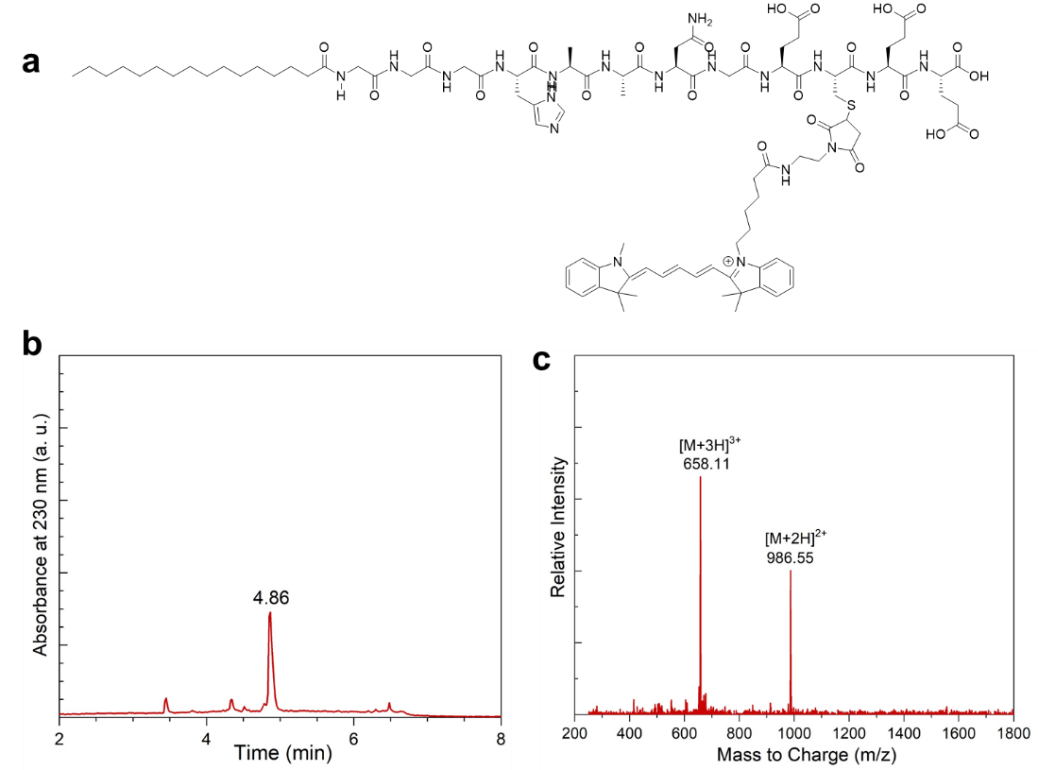

**Figure S10. Synthesis of Cy5 conjugated SA-E.** a) Molecular structure of Cy5 conjugated SA-E. b) LC and c) MS traces of the HPLC purified product.

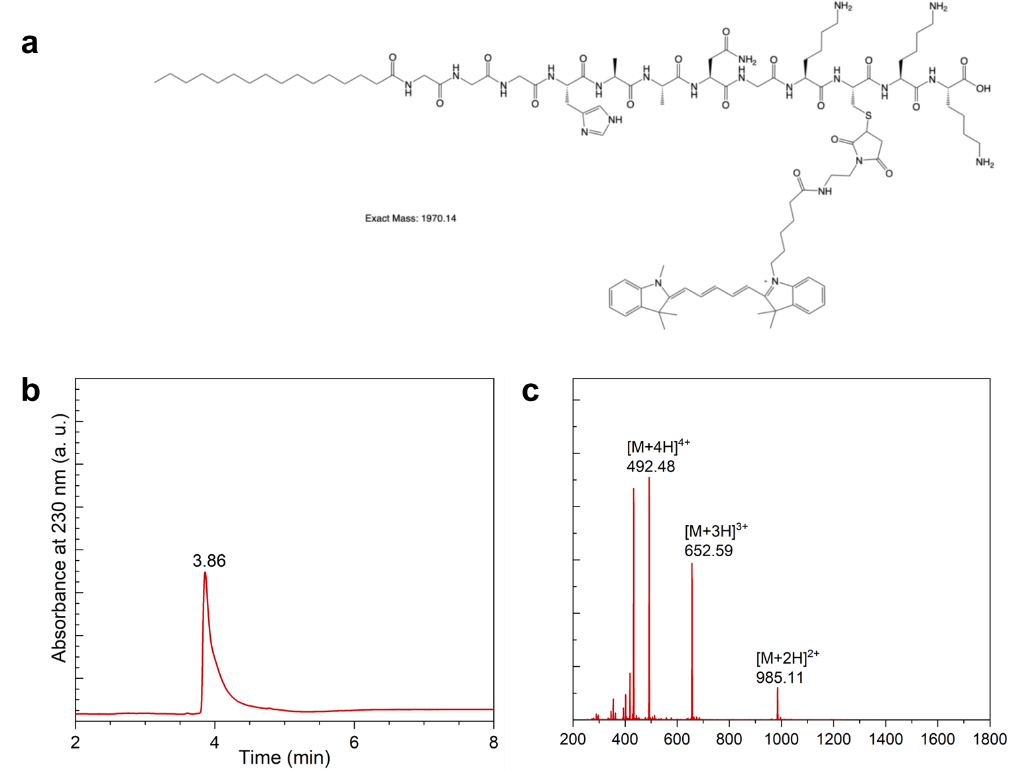

**Figure S11. Synthesis of Cy5 conjugated SA-K.** a) Molecular structure of Cy5 conjugated SA-K. b) LC and c) MS traces of the HPLC purified product.

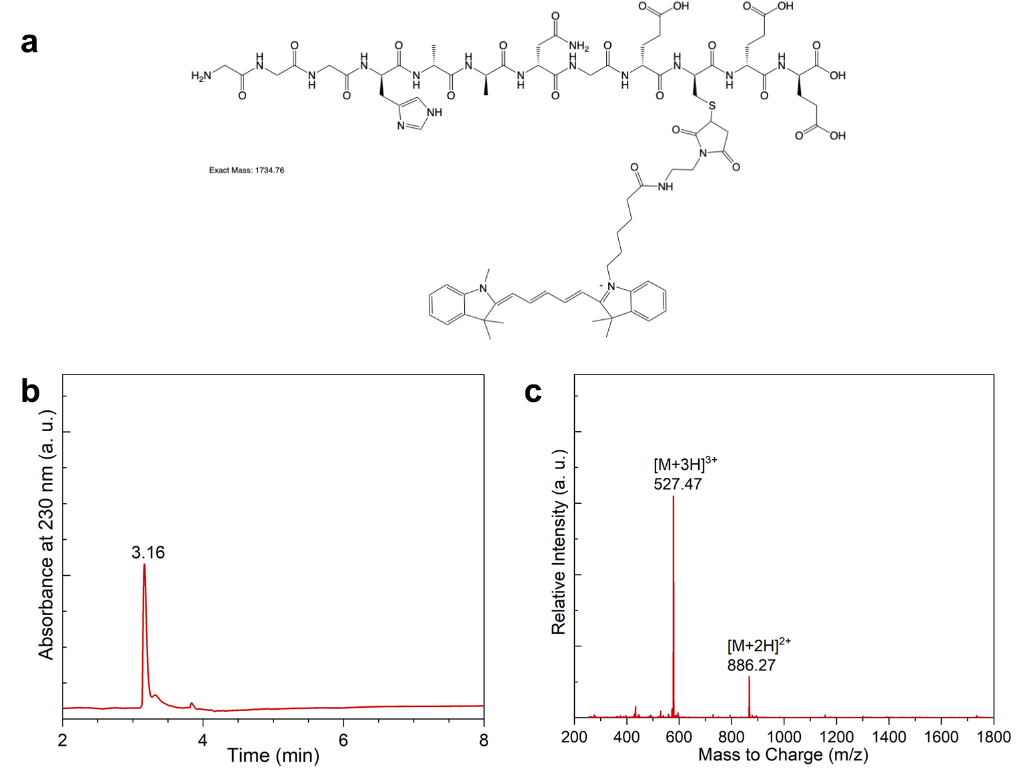

**Figure S12. Synthesis of Cy5 conjugated No-SA.** a) Molecular structure of Cy5 conjugated No-SA. b) LC and c) MS traces of the HPLC purified product.

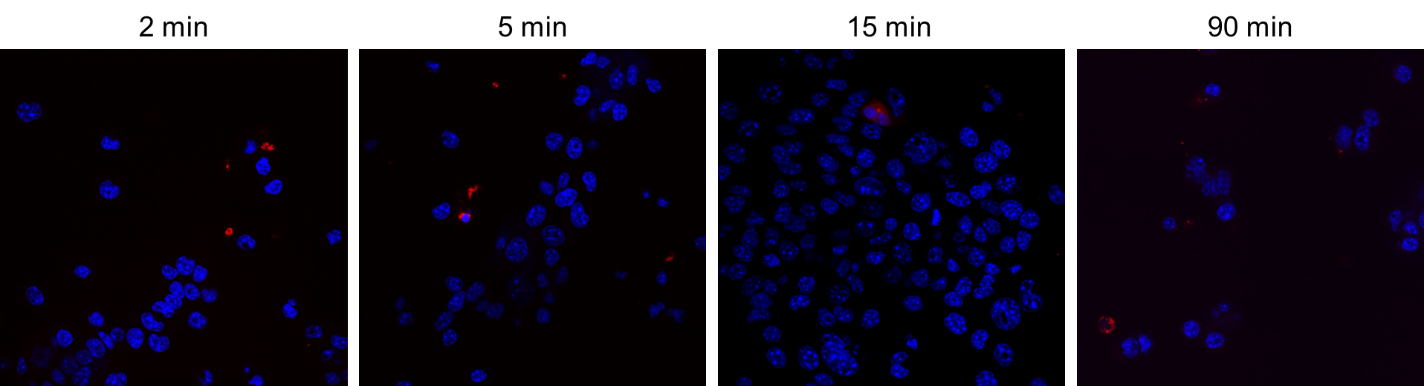

**Figure S13. No-SA was not taken up by cells.** a) Confocal microscope images of 4T1 cells incubated with No-SA at different time points. Blue: nuclei, Red: Cy5 labeled No-SA.

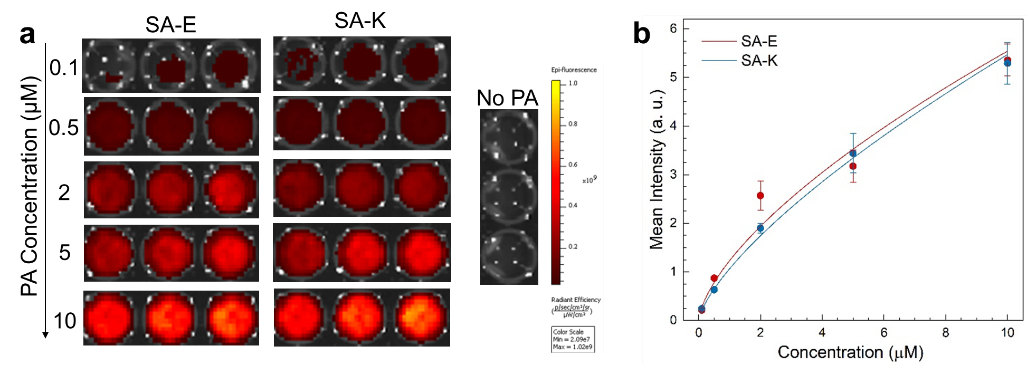

**Figure S14. Fluorescence of PAs in liver homogenates.** a) IVIS images and b) calculated mean intensities of SA-E and SA-K at different concentrations in liver homogenates. Data are presented as mean ± SEM.

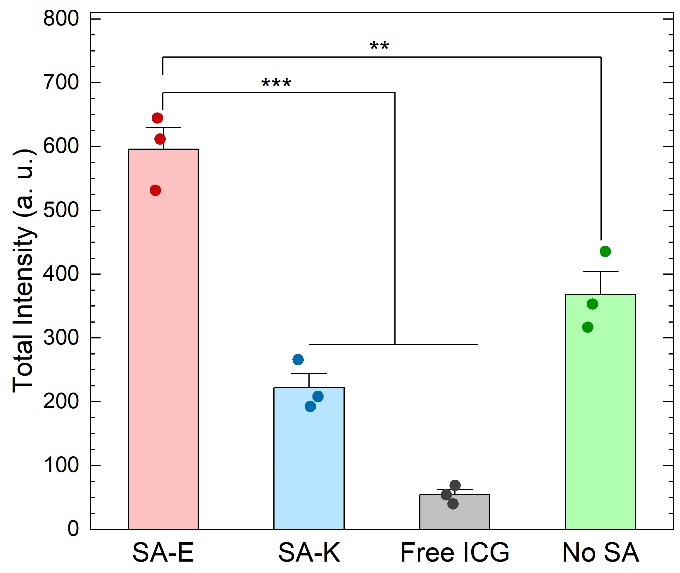

**Figure S15. Total tumor accumulation of PAs and free ICG in 4T1 tumor.** Area under the curve values of tumor signal plots for SA-E, SA-K, No-SA, and free ICG injections in Figure 3b or Figure S 16. Data are presented as mean ± SEM. Statistical analysis was performed using one-way analysis of variance (ANOVA). ***p* <0.01, and ****p* <0.001.

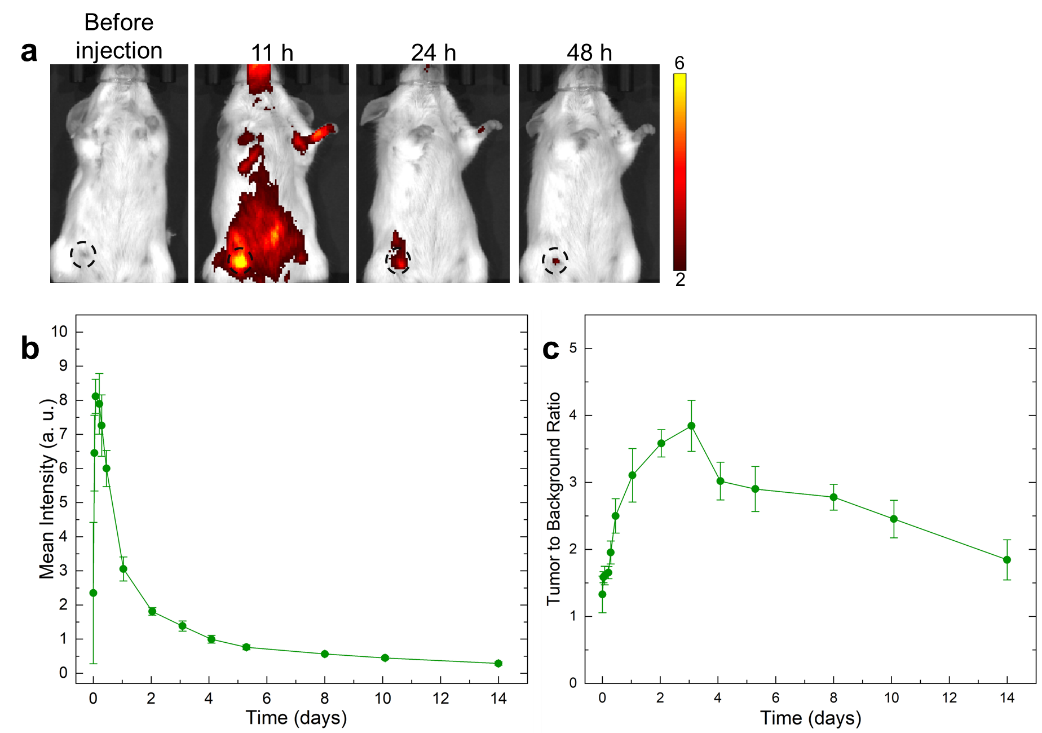

**Figure S16. Tumor accumulation of No-SA.** a) Representative IVIS images of intravenously injected ICG labeled No-SA (50 nmole) in 4T1 tumor-bearing mice at different time points. Black circles highlight the tumor location. b) Calculated mean intensities of ICG signal and c) tumor to background signal ratio of No-SA at different time points after injection. Data are presented as mean ± SEM.

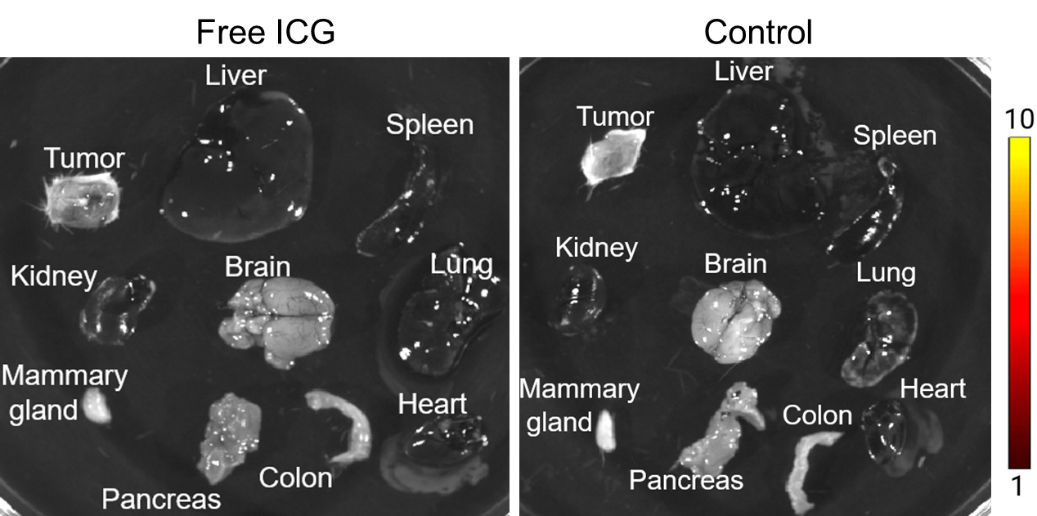

**Figure S17. Biodistribution of free ICG 2 days after injection (50 nmole).** Representative IVIS images showing free ICG signal in the tumor and other tissues were not significantly different from the control.

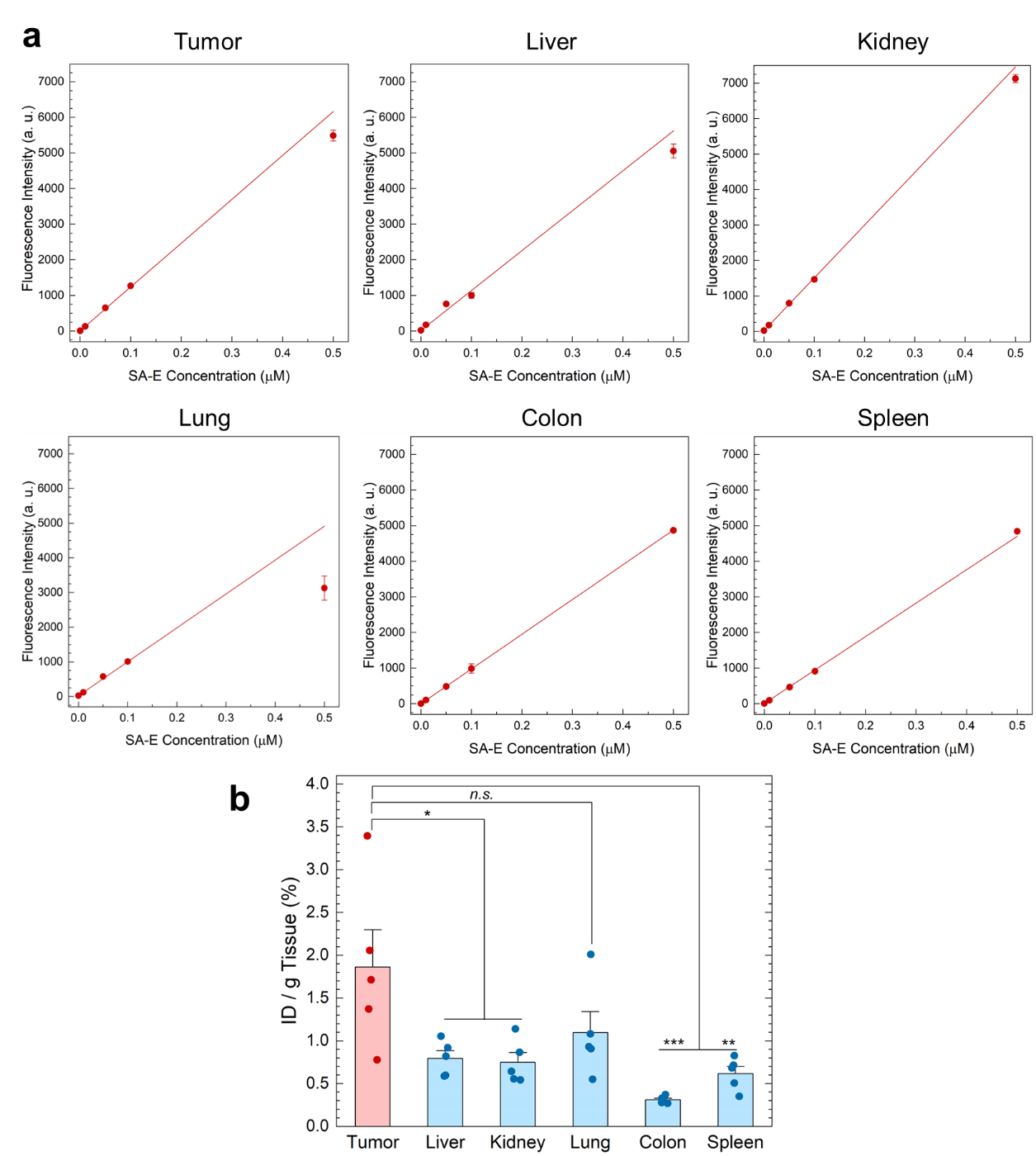

**Figure S18. Biodistribution of SA-E.** a) Calibration curves of ICG labeled SA-E fluorescence in homogenates of different tissues. b) Percent injected dose of SA-E per gram tissue in the tumor and other organs. Data are presented as mean ± SEM. Statistical analysis was performed using one-way analysis of variance (ANOVA). *n.s*. is non-significant, ***p* <0.01, and ****p* <0.001.

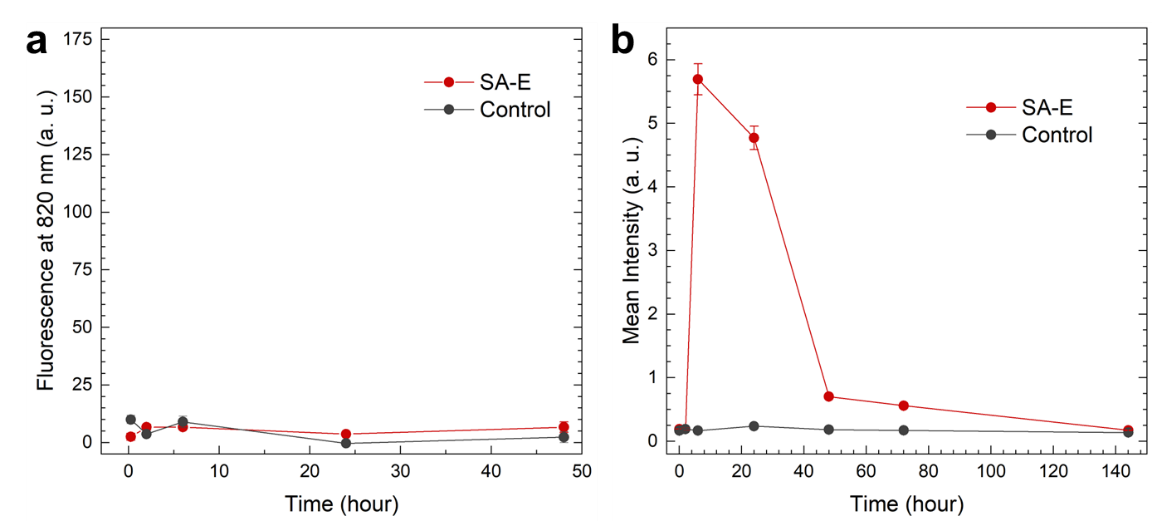

**Figure S19. SA-E was cleared through stool.** a) Fluorescence of ICG in urine samples collected from mice injected with ICG labeled SA-E (50 nmole) or PBS at different time points. Fluorescence intensity in urine samples was measured using a plate reader. c) Mean ICG intensity in stool samples collected at different time points after SA-E (50 nmole) or PBS injection measured by using an IVIS. Data are presented as mean ± SEM.

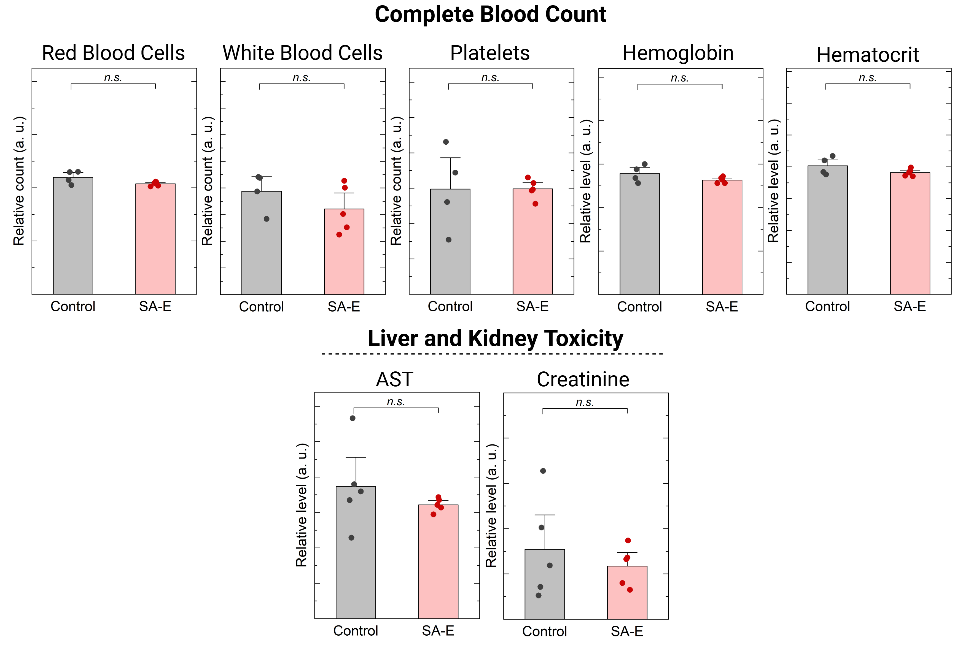

**Figure S20. Intravenously injected SA-E (200 nmole) showed no toxicity in wild type mice.** Blood toxicity counts (top panel), and liver and kidney toxicity markers (bottom panel) did not show any significant change. Control is PBS injected mice. Data are presented as mean ± SEM. Studies were run in at least triplicates. Statistical analysis was performed using Student’s t test in. *n.s.* is non-significant.

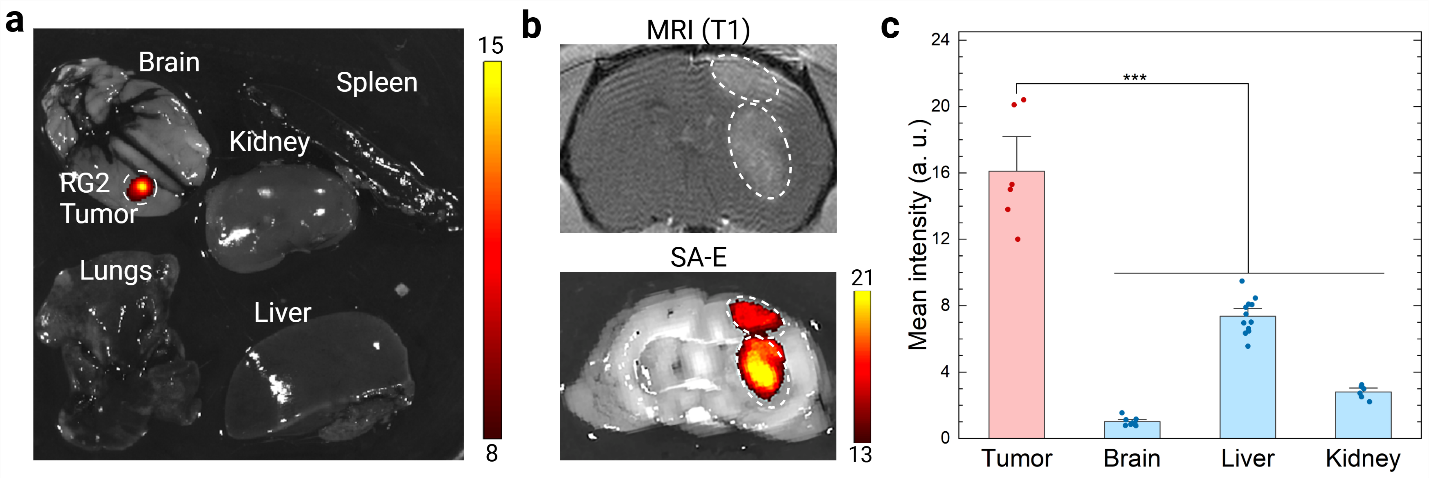

**Figure S21. SA-E demonstrated strong tumor accumulation in an intracerebral orthotopic glioma model in rats.** a) IVIS imaging of excised organs of a rat bearing a syngeneic RG2 tumor. Organs were harvested 2 days after intravenous injection of ICG labeled SA-E (500 nmole). b) Gadolinium contrast-enhanced T1 MRI and IVIS imaging of RG2 tumors. MRI imaging was performed shortly before scarifying the rats. IVIS imaging was performed using wholemount dissected brain samples. c) Accumulation of SA-E in RG2 and GBM39 tumors, healthy brain tissue, liver, and kidney in rats. Data are presented as mean ± SEM. Studies were run in at least triplicates. Statistical analysis was performed using one-way analysis of variance (ANOVA). ****p* <0.001.

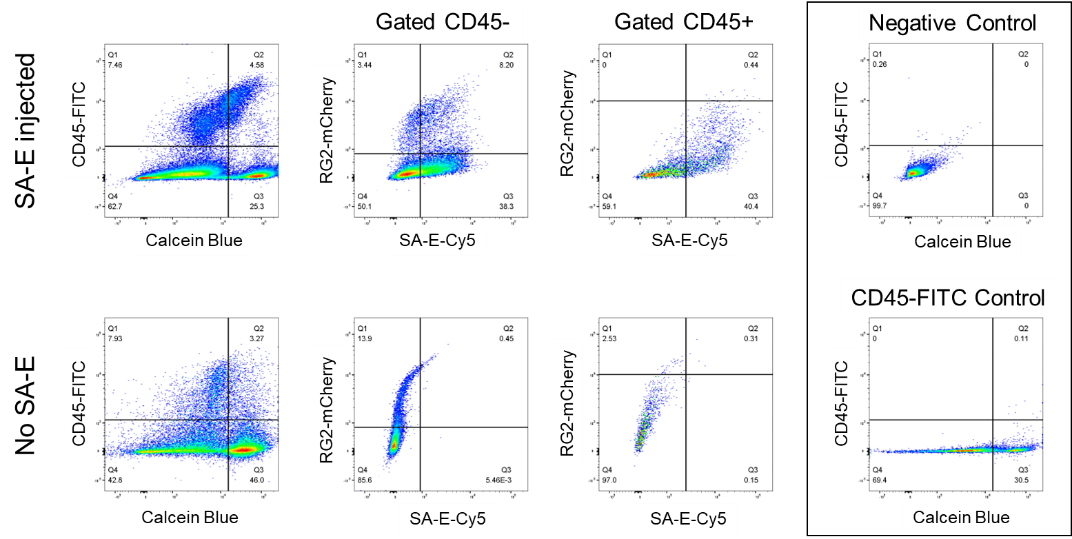

**Figure S22. Representative flow cytometry plots** showing cell populations positive for mCherry (RG2 cells), Cy5 (SA-E), and CD45 (white blood cells) in RG2 tumors with or without SA-E injection. Calcein blue was used to detect viable cell populations.

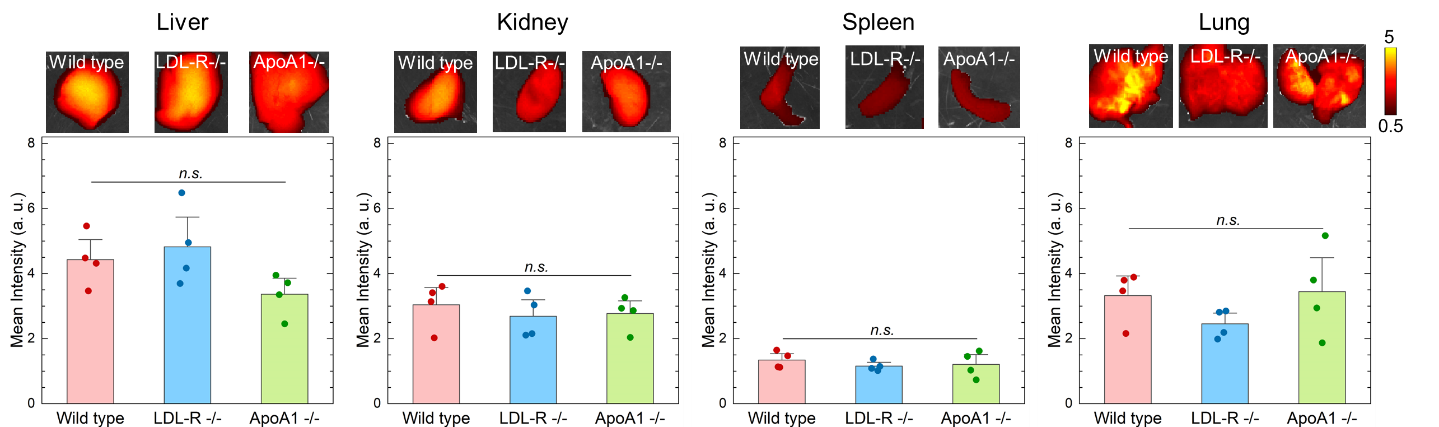

**Figure S23. Accumulation of SA-E in the major organs of wild type and knockout mice.** Mean SA-E intensity at organs collected at 2 days after injection (50 nmole). Data are presented as mean ± SEM. Statistical analysis was performed using two-way analysis of variance (ANOVA). *n.s.* is non-significant.

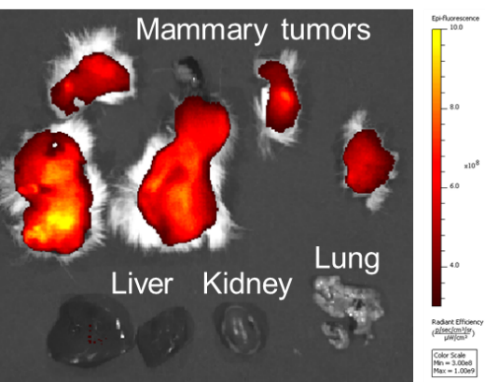

**Figure S24. Accumulation of SA-E in MMTV mice.** IVIS imaging of the harvested organs of an MMTV mouse showing specific SA-E accumulation in mammary tumors.

**Supporting Table 2.** Number of detected lesions with or without SA-E signal in the small intestines and colons of APC^min^ mice.

|  | **Small Intestine** | | | **Colon** | | |
| --- | --- | --- | --- | --- | --- | --- |
| **Mouse** | **Number of Lesions** | **SA-E Positive** | **SA-E Negative** | **Number of Lesions** | **SA-E Positive** | **SA-E Negative** |
| 1 | 15 | 15 | 0 | 2 | 2 | 0 |
| 2 | 22 | 22 | 0 | 1 | 1 | 0 |
| 3 | 34 | 34 | 0 | 1 | 1 | 0 |
| **Total** | 71 | 71 | 0 | 4 | 4 | 0 |

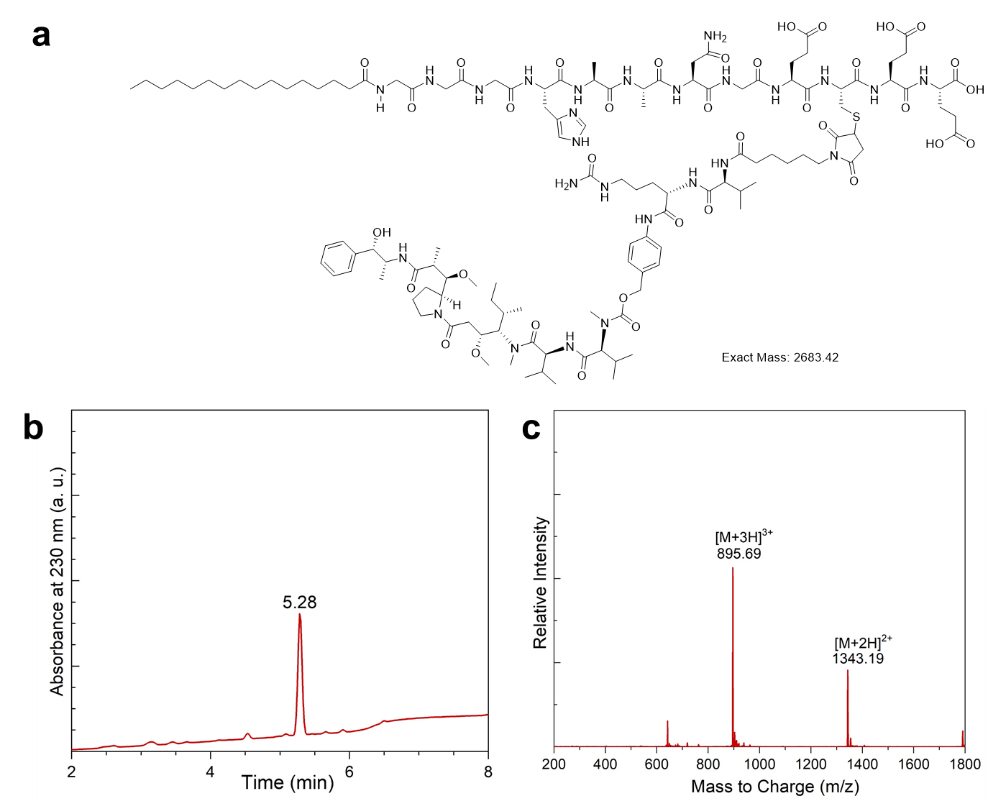

**Figure S25. Synthesis of MMAE conjugated SA-E.** a) Molecular structure of MMAE conjugated SA-E. b) LC and c) MS traces of the HPLC purified product.

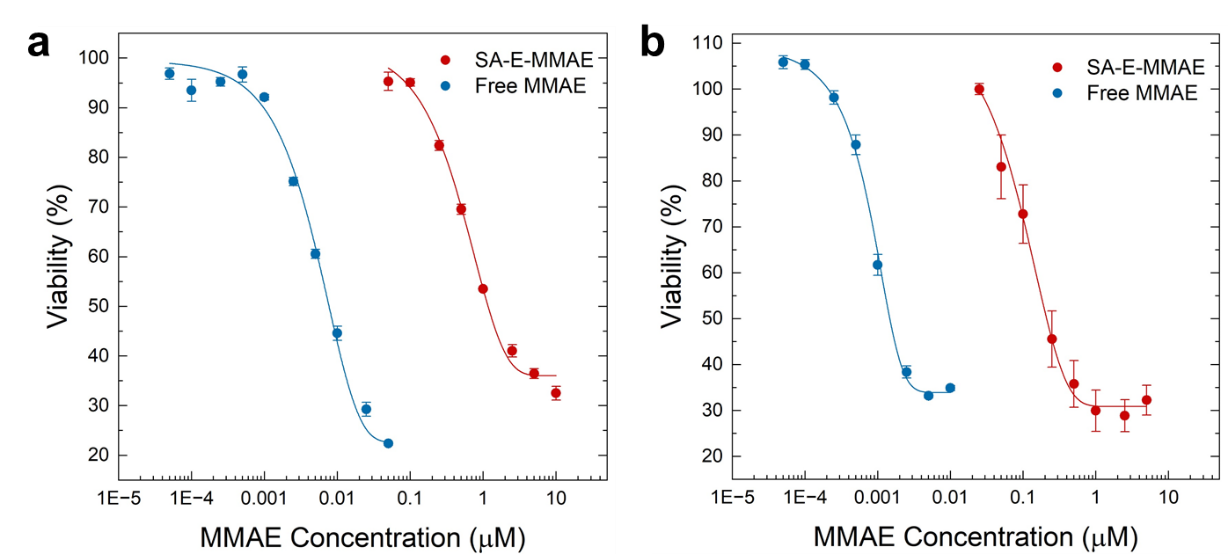

**Figure S26. Cytotoxicity of SA-E-MMAE and free MMAE against 4T1 and RG2 cells.** a) 4T1 cells and b) RG2 cells. Compared to free MMAE, SA-E-MMAE showed ~140 and ~150 fold less toxicity against 4T1 and RG2 cells with IC50 values of 1.18 µM and 0.21 µM, respectively. A decrease in the cytotoxicity of prodrugs can be expected as these drugs need to be activated by enzymes first. Data are presented as mean ± SEM.

**Supporting Methods**

**Molecular Dynamics (MD) Simulations.** We used a custom forcefield combining Amber-03 for the peptide and General AMBER Forcefield (GAFF) for the hydrophobic tails. All MD simulations were performed using Gromacs-2020 package.^1^ The simulation system consisted of the peptides in explicit solvent (water) in a 150 x 75 x 75 nm^3^ 3D periodic box. Counter ions, Na^+^ or Cl^−^, were also added to each solvated system to create neutrality. An energy minimization was performed to prevent any overlap of atoms, followed by a 1 ns equilibration run. The system was then simulated for a production run of 0.5 s. The leap-frog algorithm with a 2 fs time step was used to integrate the equations of motion. The system was maintained at 300 K and 1 bar, using the velocity rescaling thermostat^2^ and Parrinello-Rahman barostat,^3^ respectively. Particle mesh Ewald (PME)^4^ algorithm with a real space cut-off of 1.2 nm was used to calculate long-range interactions. Similarly, LJ interactions were also truncated at 1.2 nm. For the water molecules, TIP3P model^5^ was used, and LINCS^6^ algorithm was used to constrain the motion of the hydrogen atoms bonded to heavy atoms. Configurations were stored every 100 ps for visualization and analysis using PyMOL.^7^

**Mass spectroscopy analysis:**

**Samples Preparation.** Eluted plasma samples were dried down and stored at -80°C. Samples were thawed at room temperature for 10 min, and 75 µL of SDS protein extraction buffer (5% SDS, 50 mM tetraethylammonium bromide; TEAB) was added to each sample. Samples were shaken in the Thermo mixer for 10 minutes at room temperature and centrifuged at 10,000 g for 5 min. Each sample was transferred from a 0.5 mL tube to a 1.5 mL LoBind centrifuge tube. Samples were reduced by adding 3.4 µL of 0.5 M dithiothreitol and incubated at 95 °C for 10 min. Samples were alkylated by the addition of 6.8 µL of 0.5 M iodoacetamide and were incubated at room temperature for 30 min in the dark. Samples were then acidified by the addition of 8.52 µL of 12% phosphoric acid, and 562 µL of SDS protein binding buffer (90% methanol, 100 mM TEAB, pH 7.51). Samples were transferred 165 µL at a time to S-trap micro columns (Protifi, Farmingdale, NY) and inserted into 1.5 mL of polypropylene tubes. Samples were centrifuged at 4,000 g for 3 min between each addition of the sample. S-trap columns were washed 6 times using 150 µL of 90% methanol, 100 mM of TEAB followed by centrifugation at 4,000g for 3 min between each wash step. S-trap columns containing the bound sample proteins were transferred to 1.5 mL Lobind centrifuge tubes and 40 µL of 80 ng/µL sequencing grade modified trypsin (Pierce TM Trypsin Protease MS-Grade, XG348166) in 50 mM TEAB was added. S-trap columns were loosely capped, and digestion was performed at 37°C overnight in a humidified chamber.

After completion of digestion, peptides were eluted by sequential addition of 40 µL of TEAB, 40 µL of 0.2% aqueous formic acid, and 40 µl of 50% acetonitrile, 0.2% formic acid, with centrifugation at 4,000 g for 4 min between each addition of elution buffer. The elution fractions were combined and then dried by vacuum centrifugation. Samples were reconstituted by adding 20 µL of 5% formic acid, shaking at 37 °C, and spinning at 15,000 g for 5 min. The supernatant of each sample was transferred to an auto-sample vial.

**LC/MS Analysis.** 20 µL of each sample was injected into QExactive HF and run with the 90 min LC/MS method. Protein digests were separated using liquid chromatography with a Dionex RSLC UHPLC system (Thermo Fisher), then delivered to a QExactive HF (Thermo Fisher) using electrospray ionization with a Nano Flex Ion Spray Source (Thermo Fisher) fitted with a 20 µm stainless steel nano-bore emitter spray tip and a 1.0 kV source voltage. Xcalibur version 4.0 was used to control the system. Samples were applied at 10 µL/min to a Symmetry C18 trap cartridge (Waters) for 10 min, then switched onto a 75 µm x 250 mm NanoAcquity BEH 130 C18 column with 1.7 µm particles (Waters) using mobile phases water (A) and acetonitrile (B) containing 0.1% formic acid. A 7.5-30% acetonitrile gradient was applied over 60 min at a flow rate of 300 nL/min. Survey mass spectra were acquired over an m/z range of 375−1400 at 120,000 resolution (m/z 200). Data-dependent acquisition selected the top 10 most abundant precursor ions for tandem mass spectrometry by HCD fragmentation using an isolation width of 1.2 m/z, a normalized collision energy of 30, and a resolution of 30,000. Dynamic exclusion was set to auto, charge state for MS/MS +2 to +7, a maximum ion time 100 ms, a minimum AGC target of 3 x 10^6^ in MS1 mode and 5 x 10^3^ in MS2 mode.

**Data analysis.** Mass spectrometry data from all samples were processed using COMET/PAWS against Uniprot_Human_database. Comet (v. 2016.01, rev. 3)^8^ was used to search MS2 Spectra against a January 2023 version of canonical FASTA protein database containing human uniprot sequences, and concatenated sequence-reversed entries to estimate error thresholds and 179 common contaminant sequences and their reversed forms. The database processing was performed using python scripts available at https://github.com/pwilmart/fasta_utilities.git and Comet results were processed using the PAW pipeline2 from https://github.com/pwilmart/PAW_pipeline.git.

Comet searches for all samples were performed with trypsin enzyme specificity with monoisotopic parent ion mass tolerance set to 1.25 Da and monoisotopic fragment ion mass tolerance set at 1.0005 Da. A static modification of +45.9877 Da was added to all cysteine residues and a variable modification of +15.9949 Da on Methionine residues. We used a linear discriminant transformation to improve the identification sensitivity from the Comet analysis.^9,10^ Comet scores were combined into linear discriminant function scores, and discriminant score histograms created separately for each peptide charge state (2+, 3+, and 4+). Separate histograms were created for matches to forward sequences and for matches to reversed sequences for all peptides of seven amino acids or longer. The score histograms of reversed matches were used to estimate peptide false discovery rates (FDR) and set score thresholds for each peptide class. The overall protein FDR was 1.2%. Spectral counts were normalized for both sample groups.

**References**

1. Abraham, M.J. et al. GROMACS: High performance molecular simulations through multi-level parallelism from laptops to supercomputers. *SoftwareX* **1**, 19-25 (2015).

2. Bussi, G., Donadio, D. & Parrinello, M. Canonical sampling through velocity rescaling. *J. Chem. Phys*. **126**, 014101 (2007).

3. Berendsen, H.J.C., Postma, J.P.M., van Gunsteren, W.F., DiNola, A. & Haak, J. R. Molecular dynamics with coupling to an external bath. *J. Chem. Phys.* **81**, 3684-3690 (1984).

4. Darden, T., York, D. & Pedersen, L. Particle mesh Ewald - an N.LOG(N) method for Ewald sums in large systems. *J. Chem. Phys*. **98**, 10089-10092 (1993).

5. Jorgensen, W.L. et al. Comparison of simple potential functions for simulating liquid water. *J. Chem. Phys.* **79**, 926-935 (1983).

6. Hess, B., Bekker, H., Berendsen, H. & Fraaije, J. LINCS: a linear constraint solver for molecular simulations. *J. Comput. Chem.* **18**, 1463-1472, (1997).

7. PyMOL, The PyMOL Molecular Graphics System, Version 2.0 Schrödinger, LLC.

8. Eng, J.K., Jahan, T.A. & Hoopmann, M.R., Comet: an open source MS/MS sequence database search tool. *Proteomics* **13**, 22-24 (2013).

9. Wilmarth, P.A., Riviere, M.A. & David, L.L. Techniques for accurate protein identification in shotgun proteomic studies of human, mouse, bovine, and chicken lenses. *J. Ocul. Biol. Dis. Inform.* **2**, 223-234 (2009).

10. Keller, A., Nesvizhskii A.I., Kolker, E. & Aebersold, R. Empirical statistical model to estimate the accuracy of peptide identifications made by MS/MS and database search. *Anal. Chem.* **74**, 5383-5392 (2002).
